# Gene amplification acts as a molecular foothold to facilitate cross-species adaptation and evasion of multiple antiviral pathways

**DOI:** 10.1101/2022.06.06.494757

**Authors:** Shefali Banerjee, Cathy Smith, Adam Geballe, Stefan Rothenburg, Jacob O. Kitzman, Greg Brennan

**Affiliations:** Department of Medical Microbiology and Immunology, School of Medicine, University of California, Davis, Davis, CA 95616, USA, Current address for SB: Department of Biochemistry and Molecular Biology, University of Texas Medical Branch Galveston, Texas, USA; Departments of Human Genetics and Computational Medicine & Bioinformatics, University of Michigan, Ann Arbor, MI 48109, USA; Divisions of Human Biology and Clinical Research, Fred Hutchinson Cancer Center, Seattle, WA, 98109 USA; Departments of Microbiology and Medicine, University of Washington, Seattle, WA, 98195, USA

## Abstract

Cross-species spillover events are responsible for many of the pandemics in human history including COVID-19; however, the evolutionary mechanisms that enable these events are poorly understood. We have previously modeled this process using a chimeric vaccinia virus expressing the rhesus cytomegalovirus-derived PKR antagonist RhTRS1 in place of its native PKR antagonists; E3L and K3L (VACVΔEΔK+RhTRS1). Using this virus, we demonstrated that gene amplification of *rhtrs1* occurred early during experimental evolution and was sufficient to fully rescue virus replication in partially resistant African green monkey (AGM) fibroblasts. Notably, this rapid gene amplification also allowed limited virus replication in otherwise completely non-permissive human fibroblasts, suggesting that gene amplification may act as a “molecular foothold” to facilitate viral adaptation to multiple species. In this study, we demonstrate that there are multiple barriers to VACVΔEΔK+RhTRS1 replication in human cells, mediated by both PKR and RNase L. We experimentally evolved three AGM-adapted virus populations in human fibroblasts. Each population adapted to human cells bimodally, via an initial 10-fold increase in replication after only two passages followed by a second 10-fold increase in replication by passage nine. Using our Illumina-based pipeline, we found that some SNPs which had evolved during the prior AGM adaptation were rapidly lost, while 13 singlebase substitutions and short indels increased over time, including two SNPs unique to HFF adapted populations. Many of these changes were associated with components of the viral RNA polymerase, although no variant was shared between all three populations. Taken together, our results demonstrate that *rhtrs1* amplification was sufficient to increase viral tropism after passage in an “intermediate species” and subsequently enabled the virus to adopt different, species-specific adaptive mechanisms to overcome distinct barriers to viral replication in AGM and human cells.

## Introduction

Over the past 70 years, zoonotic pathogens have been responsible for more than 60% of all emerging infectious diseases in humans, including those caused by HIV, avian influenza A virus, and MERS-CoV^1^. Predicting cross-species transmission candidates before they occur is difficult because there are potentially hundreds of thousands of unknown animal viruses circulating^2^, and natural hosts often show little or no sign of infection. Adding another layer of complexity, intermediate hosts can also play a role in this process of cross-species transmission. For example, aquatic birds are the primary host for influenza A virus (IAV), but a small number of mammalian hosts can sustain IAV infection, and at least in some cases transmit the infection to humans after reassortment or other adaptive processes^3,4^. In general, intermediate hosts can enable more frequent contact with potential new hosts and drive adaptive changes that may coincidentally enhance viral replication in new hosts. However, the evolutionary mechanisms underlying these cross-species transmission events are poorly understood.

When exposed to a new virus, a potential host is protected by a robust, multi-layered series of hurdles to overcome, including physical barriers such as the skin, receptor incompatibility that may prevent viral entry, innate barriers including host restriction factors, and adaptive immune responses. However, immune responses from different hosts are not always comparable in their activity against a given virus, and host genetics plays a key role in the defining the barriers to replication for a particular host-virus pair^5^. In the case of exposure to new viruses, some host immune proteins will likely be ineffective at inhibiting a given virus, while other proteins may, by chance, recognize the new virus and inhibit its replication^6^. Thus, viruses may need to antagonize a different array of host immune proteins to productively infect new species, and virus adaptation to a new host species may result in adaptive trade-offs that elicit a fitness cost in other species^7,8^.

One of the earliest barriers to cross-species transmission is mediated by host restriction factors. These proteins recognize a variety of pathogen associated molecular patterns (PAMPs), and initiate different antiviral responses. One such PAMP, double-stranded (ds) RNA, is produced during the replication cycle of most virus families, and activates multiple host restriction factors^9^. For example, the host restriction factor protein kinase R (PKR), is activated through a process of dimerization and autophosphorylation in the presence of dsRNA. Once activated, PKR phosphorylates the alpha subunit of eukaryotic initiation factor 2 (eIF2), ultimately leading to inhibition of translation initiation, thereby preventing virus replication^10,11^. Another dsRNA-mediated antiviral response is initiated when oligoadenylate synthase (OAS) binds dsRNA and synthesizes 2’-5’-oligoadenylates (2-5A). 2-5A is a second messenger that activates RNase L, which degrades both host and viral RNA^12–14^.

We have previously modeled the process of viral adaptation to inhibit resistant PKR using a chimeric vaccinia virus expressing the rhesus cytomegalovirus (RhCMV)-derived PKR antagonist RhTRS1 instead of its native PKR antagonists, E3L and K3L (VACVΔEΔK+RhTRS1). In this system, RhTRS1 poorly inhibits African green monkey (AGM) PKR in primary fibroblasts, and it does not inhibit human or rhesus macaque PKR at all. Using experimental evolution, we demonstrated that either gene amplification of *rhtrs1* that occurred early during experimental evolution, or two individual SNPs that subsequently evolved in VACV genes, were individually sufficient to fully rescue virus replication in AGM fibroblasts^15,16^. This gene amplification also improved VACVΔEΔK+RhTRS1 replication in otherwise completely non-permissive human and rhesus fibroblasts. Recent work suggests that gene amplification may be one of the earliest adaptive responses in poxviruses to a variety of selective pressures including host restriction factors^15,17–21^. Presumably the increased gene dosage produces more protein which overwhelms the restriction factor by a mass action like effect. These observations suggested the hypothesis that gene amplification may act as a “molecular foothold” to facilitate viral transmission to multiple otherwise resistant species.

In this study we demonstrate that, while PKR was the only barrier to replication in AGM cells, both PKR and RNase L inhibited VACVΔEΔK+RhTRS1 replication in human cells. To test our “molecular foothold” hypothesis, we experimentally evolved all three previously AGM-adapted virus populations in primary human fibroblasts. Each population adapted to replicate in human cells via a bi-modal adaptation curve. During this adaptation, we observed a rapid loss of the previously-identified AGM-adaptive missense variant in the viral RNA polymerase subunit A24R, and identified 13 single-base substitutions and short indels which increase over time, including two SNPs unique to HFF adaptation. Taken together, this work suggests a model for virus adaptation through intermediate hosts, by which infection of a partially permissive species may drive relatively non-specific adaptations like gene duplication that allow the virus to spread to otherwise non-permissive hosts. Subsequently, species-specific adaptations may evolve in these new hosts, permitting the virus to establish itself in a new species.

## Material and Methods

### Cells and viruses

A549 cells, A549 PKR^-/-^ cells, A549 RNase L^-/-^ cells, A549 PKR^-/-^ RNase L^-/-^ cells (all knockout cells kindly provided by Bernard Moss^22^) and BSC40 cells (kindly provided by Stanley Riddell) were maintained in Dulbecco’s modified Eagle’s medium (DMEM; Life Technologies) supplemented with 5% fetal bovine serum (GE Healthcare) and 1% penicillin-streptomycin (Gibco). Human foreskin fibroblasts (HFF, kindly provided by Denise Galloway) were maintained in Minimum Essential Media-α (MEM-α; VWR) supplemented with 20% FBS (GE Healthcare) and 1% penicillin-streptomycin.

VACV-βg and VACVΔEΔK+RhTRS1 were constructed as described in Child, et al.^23^ and Brennan, et al.^15^, respectively.

### Experimental Evolution

The passage 8 (p8) populations of AGM-adapted viruses were established following serial passaging of the VACVΔEΔK+RhTRS1 virus in PRO1190 cells as described previously^15^. Confluent 10 cm dishes of HFF cells were initially infected with one of each of the p8 populations of AGM-adapted viruses, designated AGM-A, AGM-B, or AGM-C (MOI = 0.1). Two days post-infection, cells were collected, pelleted and suspended in DMEM+5% FBS. After three freeze/thaw cycles, virus titers were determined on BSC40 cells by titration as described below. For every subsequent round of infection, confluent 10 cm dishes of HFF cells were infected with the progeny of the previous round of replication (MOI = 0.1) and collected as described above. Human cell passaged virus populations were named to reflect their AGM cell origin, e.g. AGM-A is the founder population for HFF-A.

To generate viral DNA, confluent 10 cm plates of HFF cells were infected with each passaged virus from each timepoint (MOI = 0.01). Two days post-infection, viral genomic DNA was isolated as previously described^24^.

### Genomic analysis

Viral genome sequencing was conducted as previously described^15^. Libraries were pooled and sequenced with paired-end 150 bp reads on an Illumina HiSeq 4000 instrument. Reads were aligned to the VACVΔEΔK+RhTRS1 reference^15^ with bwa-mem^25^. Point variants and short indels were called with freebayes^26^. Copy number (averaged over viral genomes in each pool) was computed in 50-bp windows tiling the VACV genome, by taking the mean read depth divided by the average genome-wide. *Rhtrs1* copy number was estimated by dividing by the per-sample mean depth in the single copy EGFP marker gene, to account for elevated GC content (*rhtrs1:* 58.1%; EGFP: 61.6%) relative to that of the VACV genome overall (34.5%). Structural variants were called with lumpy express (v 0.2.13), and the number of supporting split and spanning read pairs supporting each variant was tallied in each sample; only events supported by both breakpoint-containing (‘split’) and spanning read pairs were retained.

### Virus titration

Virus titers were determined in BSC40 cells by 10-fold serial dilution of infected cell lysates. Two days post-infection, BSC40 cells were washed with PBS and stained with 0.1% crystal violet. All titrations were performed in biological triplicates, and each biological replicate consisted of three (Figs. 1 and 9) or two (Figs. 2, 3, 4) technical replicates each. Statistical analysis was performed using GraphPad Prism (version 9.0.2, GraphPad Software, La Jolla, CA).

**Figure 1.**
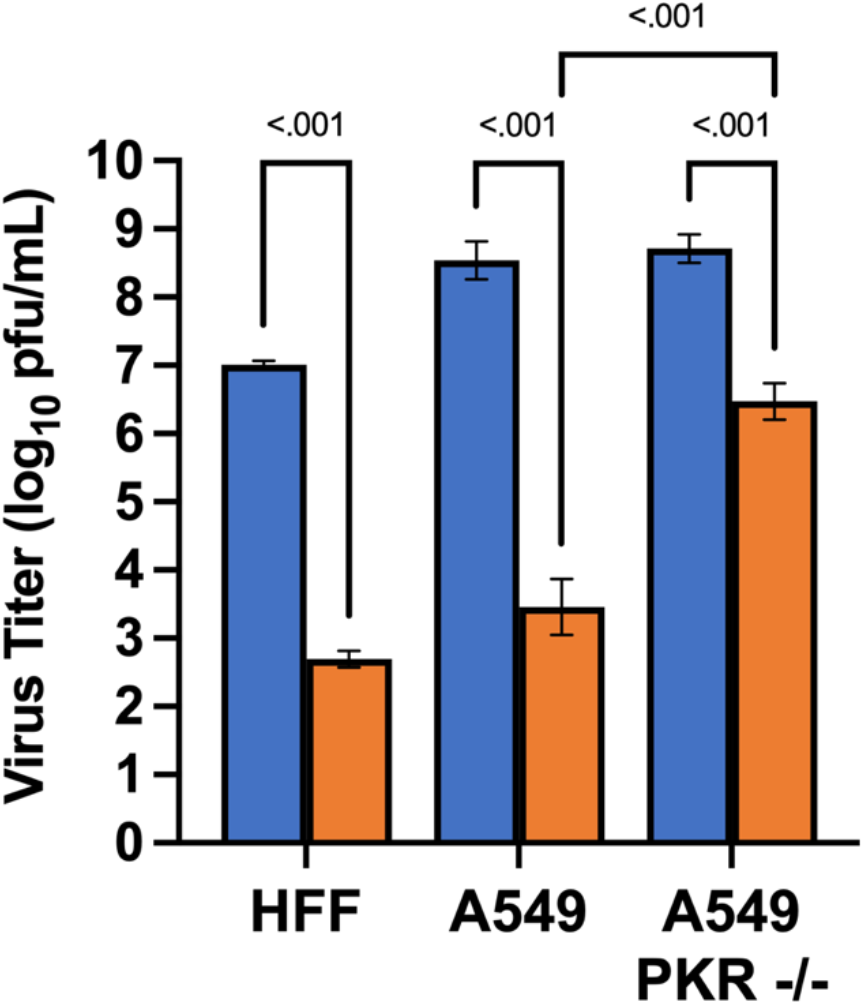
PKR knockout does not fully rescue VACVΔEΔK+RhTRS1 replication in A549 cells. Human-derived HFF, A549, <u>or A549 PKR^-/-^ cells</u> were infected with either VACV-βg or VACVΔEΔK+RhTRS1 (MOI=0.1). Two days post-infection, titers were determined by serial dilution on permissive BSC40 cells. <u>Columns represent the mean of three independent biological replicates</u>. Error bars indicate +/- one standard deviation. <u>Differences between samples were determined by multiple unpaired t-tests. Samples with adjusted p-values < 0.05 (Holm-Šídák method) are indicated by brackets with p-values indicated above</u>.

**Figure 2.**
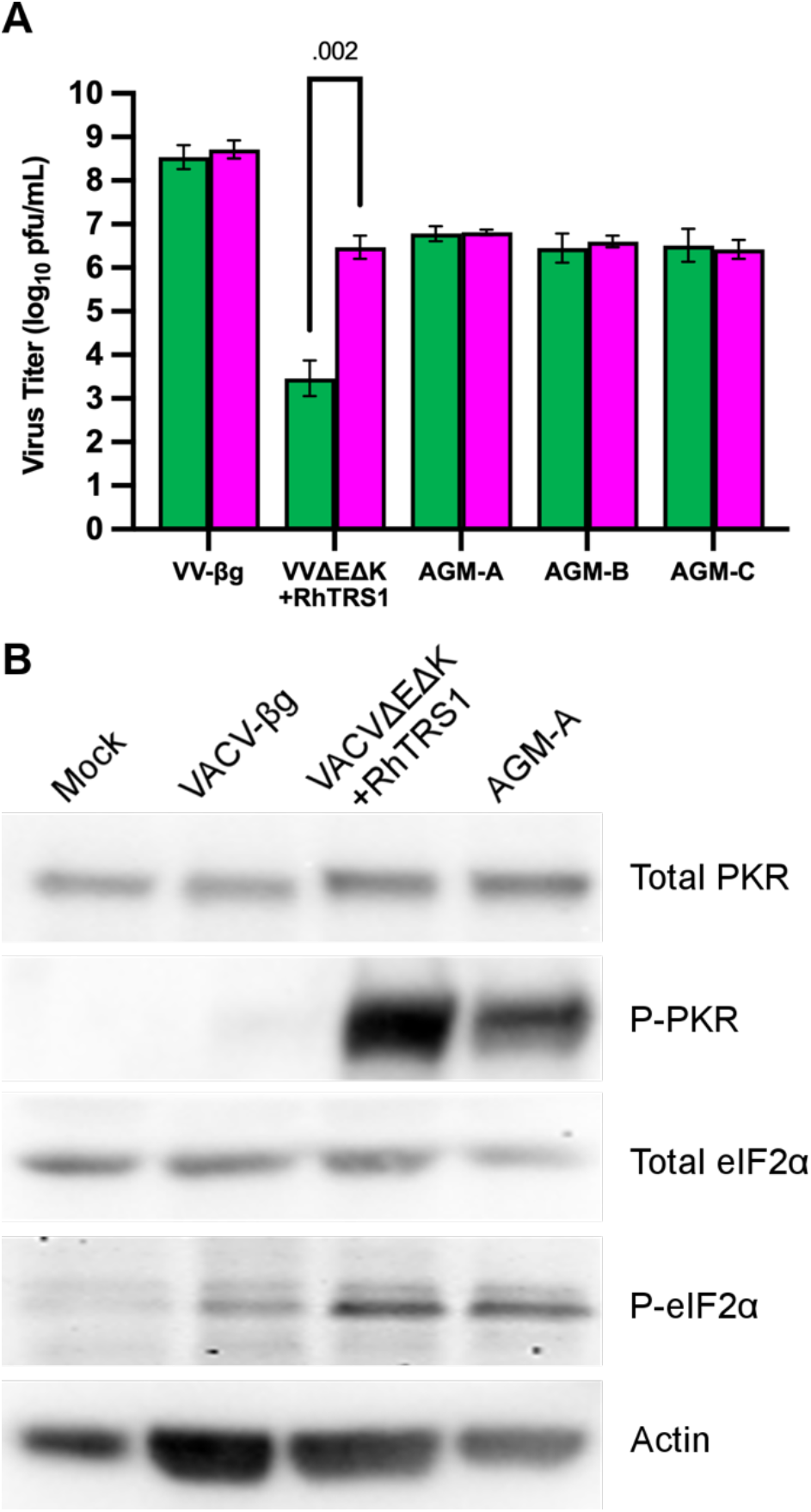
AGM-evolved SNPs and *rhtrs1* duplication do not fully rescue virus replication in human-derived cells. (A) A549 cells (green bars) or A549 PKR^-/-^ cells (magenta bars) were infected with the indicated viruses (MOI = 0.1). Two days post-infection, titers were determined by serial dilution on permissive BSC40 cells. <u>Columns represent the mean of three independent biological replicates</u>. Error bars indicate +/- one standard deviation. <u>Differences between samples were determined by multiple unpaired t-tests. Samples with adjusted p-values < 0.05 (Holm-Šídák method) are indicated by brackets with p-values indicated above</u>. (B) A549 cells were infected with the indicated viruses (MOI = 3.0). One day post-infection, cell lysates were analyzed by immunoblotting with the indicated antibodies. <u>Data are representative of three independent biological replicates</u>.

**Figure 3.**
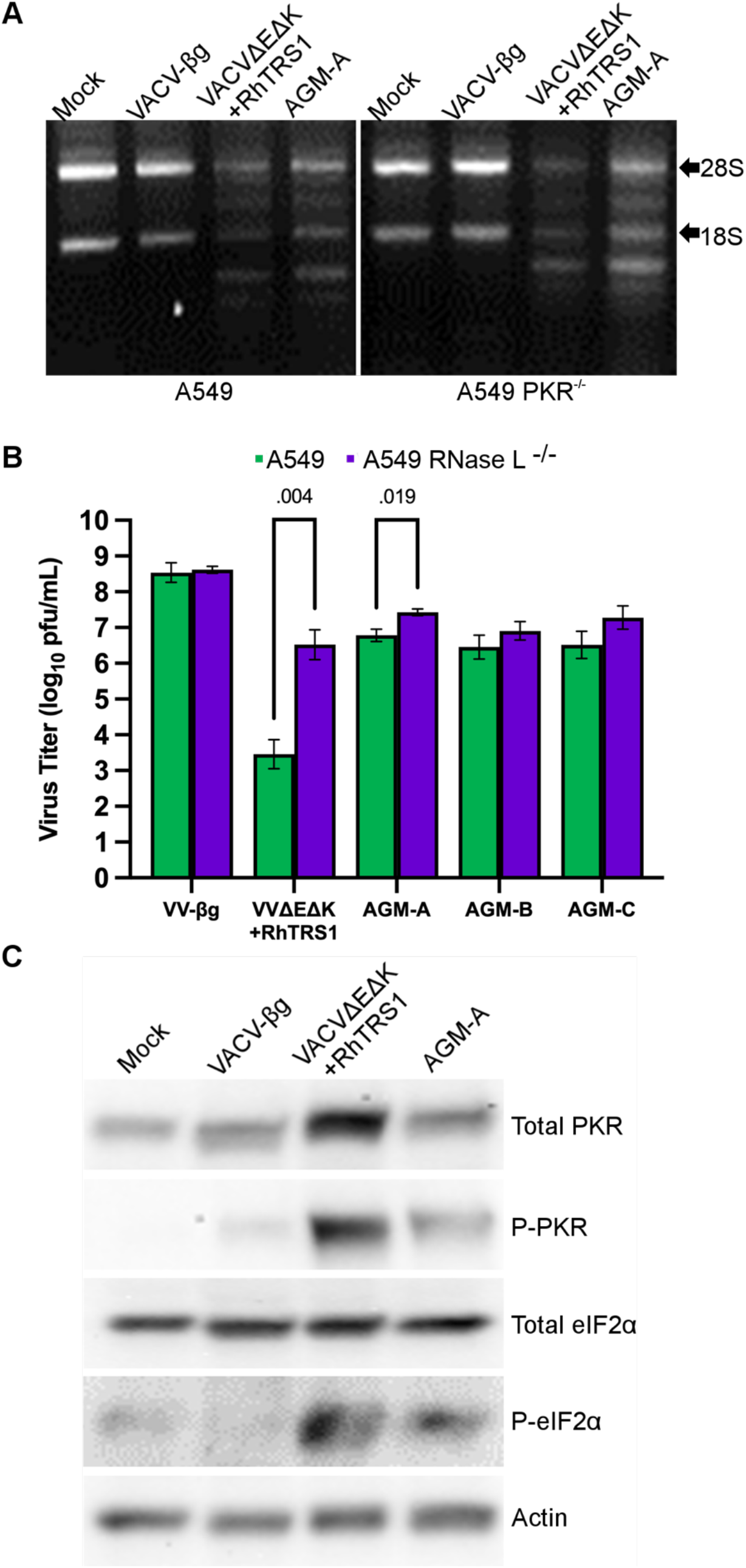
RNase L mediates a second barrier to VACVΔEΔK+RhTRS1 replication in human cells. (A) A549 cells or A549 PKR^-/-^ cells were infected with the indicated viruses (MOI = 3.0). One day post-infection, total RNA was harvested and visualized on an agarose gel + 1% bleach. (B) A549 cells (green bars) or A549 RNase L^-/-^ cells (purple bars) were infected with the indicated viruses (MOI = 0.1). Two days post-infection, titers were determined by serial dilution on permissive BSC40 cells. <u>Columns represent the mean of three independent biological replicates</u>. Error bars indicate +/- one standard deviation. <u>Differences between samples were determined by multiple unpaired t-tests. Samples with adjusted p-values < 0.05 (Holm-Šídák method) are indicated by brackets with p-values indicated above</u>. (C) A549 RNase L^-/-^ cells were infected with the indicated viruses (MOI = 3.0). One day post-infection, cell lysates were analyzed by immunoblotting with the indicated antibodies. <u>Data are representative of two independent biological replicates</u>.

### Immunoblot assay

All cells were either mock-infected or infected with the indicated viruses (MOI = 3.0). One day post-infection, the cells were lysed in 1% sodium dodecyl sulfate (SDS). Equivalent lysate volumes were separated on 10% SDS-polyacrylamide gels, then transferred to polyvinylidene difluoride (PVDF) membranes. Membranes were blocked with 5% non-fat milk dissolved in TBST (20M Tris, 150mM NaCl, 0.1% Tween 20, pH 7.4) for 1 hour and probed with one of the following primary antibodies: anti-PKR (sc-6282; Santa Cruz Biotechnology, Inc.), anti-phospho-PKR (ab32036; Abcam), anti-eIF2α or anti-phospho-eIF2α (Ser51) antibody (9722 and 9721, respectively; Cell Signaling Technology), anti-TRS1 999^27^, or anti-actin (A2066; Sigma). All primary antibodies were diluted in TBST containing 5% BSA and incubated overnight at 4°C with the membrane. Membranes were washed with TBST three times for 5 mins and then incubated for 1 hour at room temperature with either donkey anti-rabbit or goat anti-mouse secondary antibodies conjugated to horseradish peroxidase (A16110 or 62–6520, respectively; Invitrogen) at 1:10,000 in TBST containing 5% (w/v) nonfat milk. Proteins were detected using the Amersham chemiluminescent detection system (GE Healthcare) according to the manufacturer’s recommendations. Membranes were imaged using the iBright Imaging System (Invitrogen).

### RNA degradation assay

All cells were mock-infected or infected with the indicated viruses (MOI = 3.0). One day postinfection, cell lysates were harvested in TRI Reagent (Sigma). Total RNA was isolated using the Direct-zol RNA Miniprep kit (Zymo Research) following the manufacturer’s protocol. The isolated RNA was visualized using agarose gel electrophoresis as previously described^28^. Briefly, 1 μg of total RNA was loaded onto a 2% agarose gel supplemented with 1% bleach (Clorox) and electrophoresed at 50 V for 1 hour, then imaged on an iBright Imaging System (Invitrogen).

## Results

### PKR-mediated inhibition does not account for the entire replication block in human cells

We previously demonstrated cell-line specific differences in the replication of VACVΔEΔK+RhTRS1. This virus replicated well in AGM-derived BSC40 cells; however, viral replication was reduced approximately 100-fold in AGM primary fibroblasts. In addition, human primary fibroblasts (HFF) were completely resistant to VACVΔEΔK+RhTRS1 infection^15^. To determine whether immortalized human cell lines would support virus replication similar to BSC40 cells, we infected A549 cells with either VACV strain Copenhagen expressing a β-galactosidase reporter gene from the TK locus (VACV-βg)^23^, or VACVΔEΔK+RhTRS1. Unlike immortalized AGM cells, VACVΔEΔK+RhTRS1 replication was restricted in A549 cells 10,000-fold relative to VACV-βg replication, similar to the reduction we previously reported in HF cells (Fig. 1). Because A549 cells replicated the virus-resistant phenotype, we infected an existing A549 PKR knockout cell line^22^ (A549 PKR^-/-^). Similar to our previous knockdown results in HFF, eliminating PKR in A549 PKR^-/-^ cells improved VACVΔEΔK+RhTRS1 replication approximately 1000-fold relative to the PKR-competent cells (Fig. 1). However, VACVΔEΔK+RhTRS1 still replicated approximately 100-fold lower than the wild type virus in PKR^-/-^ cells.

We have previously shown that *rhtrs1* duplication provided a partial replication benefit in HFF cells, similar to the effect of PKR knockdown^15^. Therefore, we asked whether this phenotype was conserved in other human-derived cells. We infected A549 cells with VACV-βg, VACVΔEΔK+RhTRS1, or the three AGM-adapted populations, AGM-A, AGM-B, or AGM-C. All three of these populations contain an amplification of the *rhtrs1* locus and unique arrays of SNPs at various allelic frequencies in the different populations. The AGM-adapted viruses all replicated approximately 100-to 1000-fold better than VACVΔEΔK+RhTRS1, but still 100- to 1000-fold less well than VACV-βg in A549 cells (Fig. 2A, green bars), consistent with our previous observations in primary human fibroblasts.

Because neither PKR knockout nor *rhtrs1* amplification individually rescued VACVΔEΔK+RhTRS1 replication completely, we asked whether *rhtrs1* amplification would further enhance virus replication in the absence of PKR. We infected both PKR-competent and PKR^-/-^ A549 cells with all three AGM-adapted virus populations and compared the titers with VACV-βg and VACVΔEΔK+RhTRS1 infected cells. PKR knockout did not provide an additional replication benefit to any of the three AGM-adapted viruses, as they still replicated to titers approximately 100-fold lower than VACV-βg infected cells (Fig. 2A, magenta bars). These results suggested the hypothesis that *rhtrs1* amplification may be fully inhibiting PKR but there is a second block to VACVΔEΔK+RhTRS1 replication in human cells. To test this hypothesis, we performed immunoblot analysis on PKR pathway intermediates in infected A549 cells. We had limited virus stock of some populations; therefore, to avoid passaging these viruses again we performed these experiments exclusively with AGM-A to represent the AGM-adapted viruses. As expected, VACV-βg infected cells showed no PKR phosphorylation and minimal eIF2α phosphorylation, while VACVΔEΔK+RhTRS1 had much higher levels of phosphorylation of both PKR and eIF2α (Fig. 2B). PKR was also phosphorylated in AGM-A infected cells, consistent with our previous results and the published mechanism for RhTRS1-mediated inhibition of PKR^23^. However, AGM-A infected A549 cells had reduced but detectable eIF2α phosphorylation, suggesting that AGM-adapted viruses do not fully inhibit human PKR even though, paradoxically, PKR knockout does not improve AGM-A replication. Taken together, these results suggest that PKR is not the only host factor restricting VACVΔEΔK+RhTRS1 replication in human cells.

### RNase L mediates a second block to VACVΔEΔK+RhTRS1 in human cells

Since it was recently shown that RNase L cleavage products can act as PKR substrates to enhance PKR activation^29^, we asked whether RNase L also played a role in restricting VACVΔEΔK+RhTRS1 replication in human cells. To address this question, we infected A549 or A549 PKR^-/-^ cells with VACV-βg, VACVΔEΔK+RhTRS1 or AGM-A and assayed cells for RNA degradation products 24 hours post-infection. As expected, mock infected or VACV-βg infected cells did not show any evidence of RNase L activation in either cell line. However, infection with either VACVΔEΔK+RhTRS1 or the AGM-A virus population yielded multiple RNA degradation products in both A549 cells (Fig. 3A, left panel) and A549 PKR^-/-^ cells (Fig. 3a, right panel), consistent with RNase L-mediated inhibition of these viruses.

We then infected either RNase L-competent A549 cells or an existing A549 cell line containing a CRISPR-mediated deletion of RNase L^22^ (A549 RNase L^-/-^) with VACV-βg, VACVΔEΔK+RhTRS1 or the three AGM-adapted virus populations (Fig. 3B). The presence or absence of RNase L had no effect on VACV-βg replication. However, VACVΔEΔK+RhTRS1 replicated approximately 1000-fold higher in A549 RNase L^-/-^ cells relative to A549 cells, but still approximately 100-fold lower than VACV-βg titers in these cells (Fig. 3B). This increase in titer is comparable to the increase in titer we observed in A549 PKR^-/-^ cells. Unlike our data in A549 PKR^-/-^ cells, the AGM-adapted viruses also replicated somewhat better in A549 RNase L^-/-^ cells, although the magnitude of this increase was smaller than for VACVΔEΔK+RhTRS1 and no virus was fully rescued by RNase L knockout. We then performed immunoblot analysis to determine whether the PKR pathway was activated in infected A549 RNase L^-/-^ cells. Both VACVΔEΔK+RhTRS1 and AGM-A infected cells still phosphorylated both PKR and eIF2α; however, the phosphorylation of each protein was reduced in AGM-A infected cells relative to VACVΔEΔK+RhTRS1 infected cells (Fig. 3C). These data demonstrate that RNase L mediates a second, independent block to VACVΔEΔK+RhTRS1 replication in human cells.

Somewhat unexpectedly, knocking out either PKR or RNase L in A549 cells did not completely rescue replication of AGM-adapted virus populations. Combined with the immunoblot and RNA degradation data, these results support the hypothesis that *rhtrs1* duplication provides a partial replication benefit in the face of either restriction factor, without completely inhibiting either one. To test this hypothesis, we infected A549 cells carrying a CRISPR-mediated knockout of both PKR and RNase L^22^ (A549 PKR^-/-^ RNase L^-/-^) (Fig. 4). This double knockout again had no effect on VACV-βg replication; however, knocking out both PKR and RNase L improved the replication of VACVΔEΔK+RhTRS1 and all three AGM-adapted viruses to levels equivalent to VACV-βg. Taken together, these data indicate that both PKR and RNase L restrict virus replication in human cells. Furthermore, while AGM adaptation provided some resistance to both human host restriction factors, neither *rhtrs1* amplification nor the pre-existing SNPs were sufficient to fully inhibit either protein.

**Figure 4.**
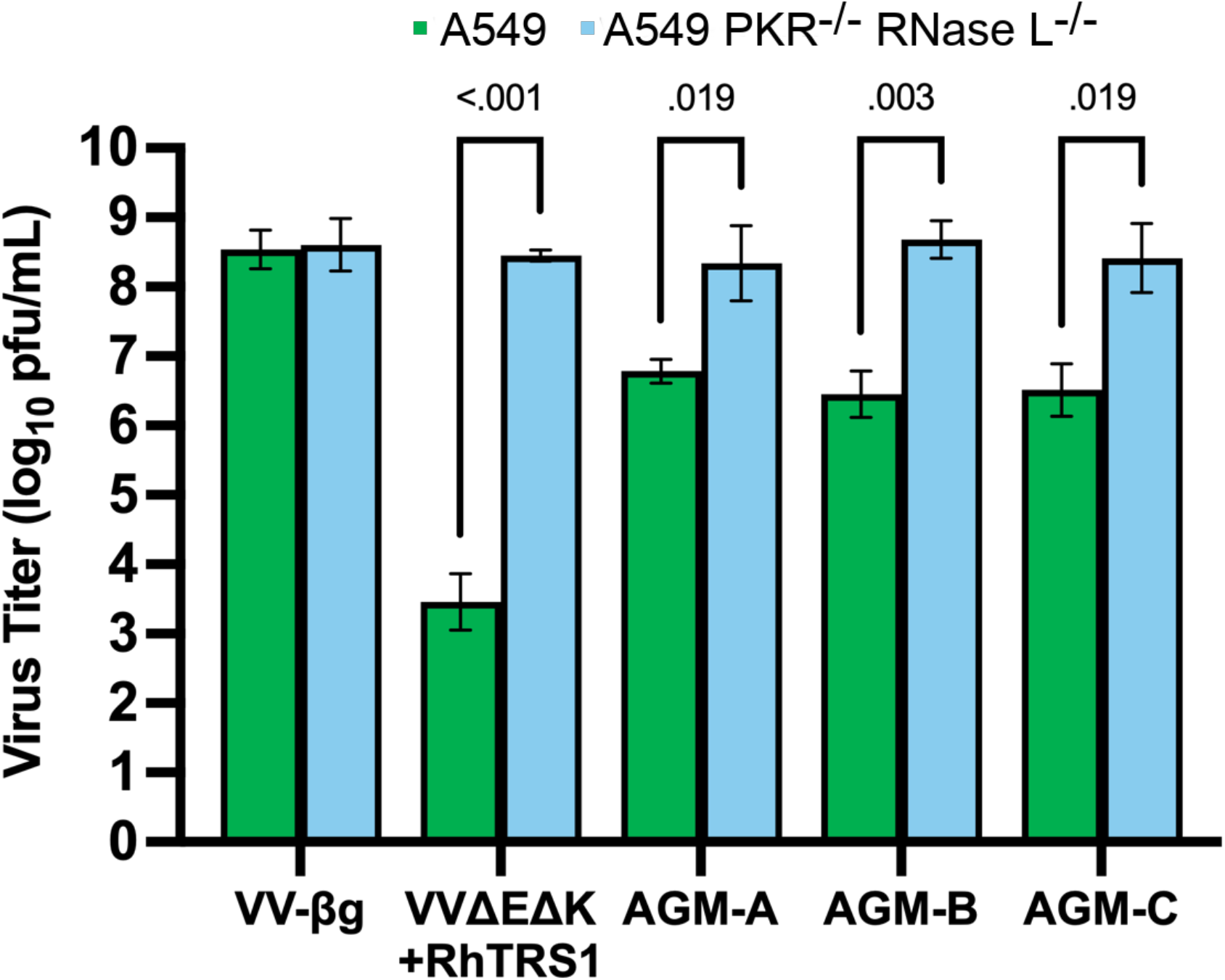
Knocking out both PKR and RNase L fully restores VACVΔEΔK+RhTRS1 replication in human cells. A549 cells (green bars) or A549 PKR^-/-^ RNase L^-/-^ cells (blue bars) were infected with the indicated viruses (MOI = 0.1). Two days post-infection, titers were determined by serial dilution on permissive BSC40 cells. <u>Columns represent the mean of three independent biological replicates</u>. Error bars indicate +/- one standard deviation. <u>Differences between samples were determined by multiple unpaired t-tests. Samples with adjusted p-values < 0.05 (Holm-Šídák method) are indicated by brackets with p-values indicated above</u>.

### *RΛtrs7*-duplicated viruses adapt to human cells in a bimodal fashion during experimental evolution

In our previous work, experimental evolution of VACVΔEΔK+RhTRS1 in HFF failed, and we were never able to detect viable virus beyond three serial passages in these cells^15^. However, because the AGM-adapted viruses did provide at least some increased fitness in human cells, we hypothesized that mutations acquired during adaptation in AGM cells may act as a “molecular foothold”, providing modest improvements to viral replication, enabling these viruses to adapt to and productively infect otherwise completely resistant human cells. To test this hypothesis, we experimentally evolved each AGM-adapted virus population independently, serially passaging these viruses in primary HFF cells at a low multiplicity of infection (MOI = 0.1). We maintained the same letter designation for each population, e.g. AGM-A is the founder population for HFF-A. For each round of serial passage, we lysed the infected cells two days post-infection (dpi), titered the resulting virus, and infected new HFF cells at the same MOI.

After only two rounds of serial passage we observed a 10-fold increase in titers for all three virus populations (HFF-Ap2, HFF-Bp2 and HFF-Cp2) (Fig. 5). This initial increase in replication was essentially stable for six rounds of serial passage. By the seventh serial passage each population started a second, more gradual 10-fold increase in virus titer. This second increase improved virus replication to titers similar to VACV-βg levels by passage 8 for the HFF-A population and by passage 9 for the HFF-B and HFF-C populations. This overall ~100-fold increase in virus replication remained stable through passage 12 in all three virus populations (Fig. 5).

**Figure 5.**
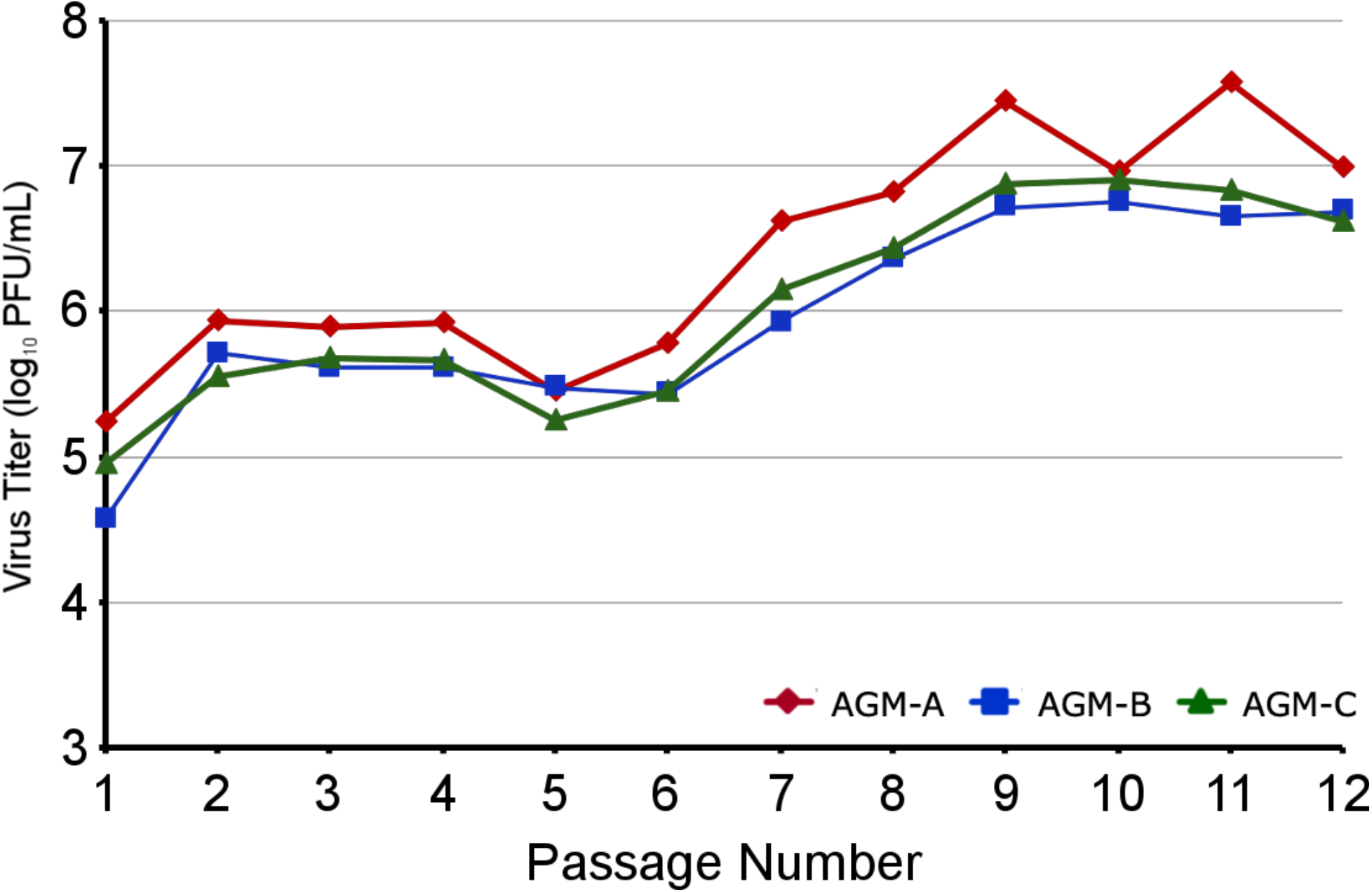
Experimental evolution of AGM-adapted viruses enables adaptation to human cells in a bimodal fashion. HFF cells were initially infected with AGM-adapted VACVΔEΔK+RhTRS1 (passage 8, previously described in ^15^ (MOI = 0.1). The A, B, C naming scheme was kept, e.g. AGM-A was the founder for HFF-A. Virus lysate was harvested two days post-infection, titered on BSC40 cells, and used to infect fresh HFF cells and the process was repeated. Three independent infections resulted in an ~ 10-fold gain of replication fitness in HFF that was evident by passage two, and a ~100-fold gain of replication fitness, evident by passages seven to nine and stable to passage 12.

To identify the genetic changes responsible for the gain in replication fitness, we used our established Illumina-based pipeline to sequence genomic DNA from each passage for all three virus populations. The sole exception to this sequencing time-course was the first passage of HFF-B, because we were unable to isolate a sufficient amount of genomic DNA from the limited sample remaining after passage. Using read depth and discordant mapping orientation as independent signals of structural variation including gene amplification, we identified only the region surrounding the *rhtrs1* locus as amplified (Fig. 6A, Table S1). In this region, we identified the same amplifications in the HFF-adapted viruses that we previously described for the AGM-adapted populations^15^. The predominant duplication shared between all three populations was 4.7 kb, spanning from L5R, upstream of the *rhtrs1* cassette, to the 3’ half of J2R, downstream of the *rhtrs1* cassette. This duplication was supported by ‘split reads’ (in which a sequenced read crosses the duplication breakpoint) and ‘spanning’ read pairs (which straddle the breakpoint). A smaller 3.4 kb duplication spanning the NeoR gene, which is part of the inserted *rhtrs1* cassette but upstream of *rhtrs1*, to the 3’ half of J2R was identified at low frequency in population HFF-A, but this shorter duplication was rapidly outcompeted during serial passage in HFF cells (Fig. 6A and Fig. S3).

**Figure 6.**
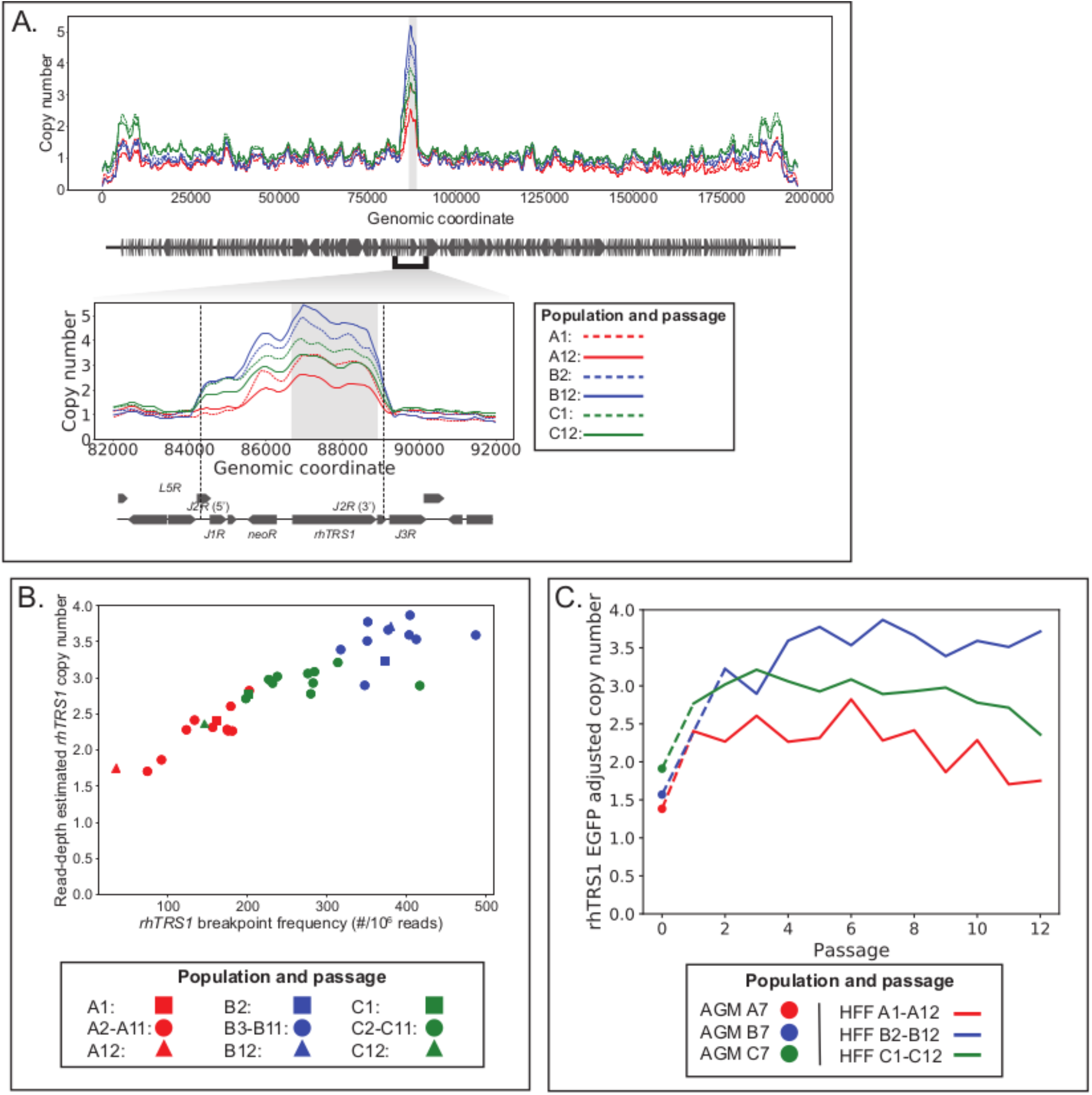
Only the *rhtrs1* locus is amplified after adaptation to human fibroblasts, and copy number increased rapidly during passage in human cells. A. Read-depth based estimate of copy number is plotted by genomic position for the first passage (dashed lines) and last passage (solid line) for each evolution across the VACV genome (*rhtrs1* region inset). Copy number was computed within 50 bp windows and then normalized by the read depth at the EGFP marker to adjust for GC content. B. Scatter plot showing correlation of read-depth based copy number estimates (y-axis) and breakpoint-containing “split reads” (x-axis) for the predominant *rhts1* duplication. First, middle, and last passages are shown as squares, circles, and triangles. C. *rhtrs1* copy number is plotted by passage for evolutions A (red), B (blue), and C (green).

To identify the dynamics of *rhtrs1* gain and loss across experimental evolution, we estimated its copy number following each round of serial passage. In all three populations at each timepoint the read-depth estimated *rhtrs1* copy number was correlated with the breakpoint frequency (Fig. 6B). We identified a rapid and early increase in *rhtrs1* copy number from the initial copy number in AGM cells for each population that correlated with the initial 10-fold increase in viral replication (compare Fig. 5 and Fig. 6C). In HFF-A, the average *rhtrs1* copy number increased from ~1.4 copies in the founder population to ~2.8 copies at passage 6. This increase was maintained until passage six, and then declined to an average of ~1.8 copies of *rhtrs1* by passage 12 (Fig. 6C, red line). A similar trend was observed for HFF-C, where the early passages increased from an initial average of ~1.9 copies to ~3.2 copies of *rhtrs1*. This number began to decline after passage nine, and by passage 12 the HFF-C population contained an average of approximately ~2.4 copies of RhTRS1 (Fig. 6C, green line). HFF-B also underwent an initial amplification from ~1.6 copies to ~3.9 copies. However, unlike HFF-A and HFF-C, we did not observe a decrease in average *rhtrs1* copy number in HFF-B (Fig. 6C, blue line).

We next performed immunoblot analysis to determine whether the increased genomic copy number correlated with an increase in RhTRS1 expression (Fig. S4). In A549 cells, VACVΔEΔK+RhTRS1 infection resulted in very little RhTRS1, while cells infected with the AGM-A virus had increased RhTRS1 expression. We did not detect RhTRS1 expression in two independent infections with HFF-Ap5 even though *rhtrs1* copy number was increased relative to AGM-A. However, HFF-Ap9 and HFF-Ap12 infected cells both had much higher RhTRS1 expression than AGM-A. HFF-Bp5- and HFF-Cp5-infected cells expressed high levels of RhTRS1. Both HFF-A- and HFF-C-infected cells had peak RhTRS1 expression at passage 9 that decreased by passage 12. HFF-B-infected cells had peak RhTRS1 expression at passage 5 that decreased but was still detectable in passages 9 and 12. Notably, both HFF-Bp12 and HFF-Cp12 infected cells expressed lower amounts of RhTRS1 than AGM-A, despite a higher *rhtrs1* copy number.

Just as in the AGM-adapted populations, we did not detect a single SNP in the *rhtrs1* locus, despite the fact that amplification of this locus provided a partial replication benefit to these viruses. Instead, we identified 26 point or short indel mutations in VACV genes which reached allelic frequencies ≥5% in at least one of the sequenced virus populations (Table 1, Table S2, and Fig. S5). Of these, 11 were pre-existing in the AGM-adapted populations, at allelic frequencies ranging from 3-73%.

**Table 1.**
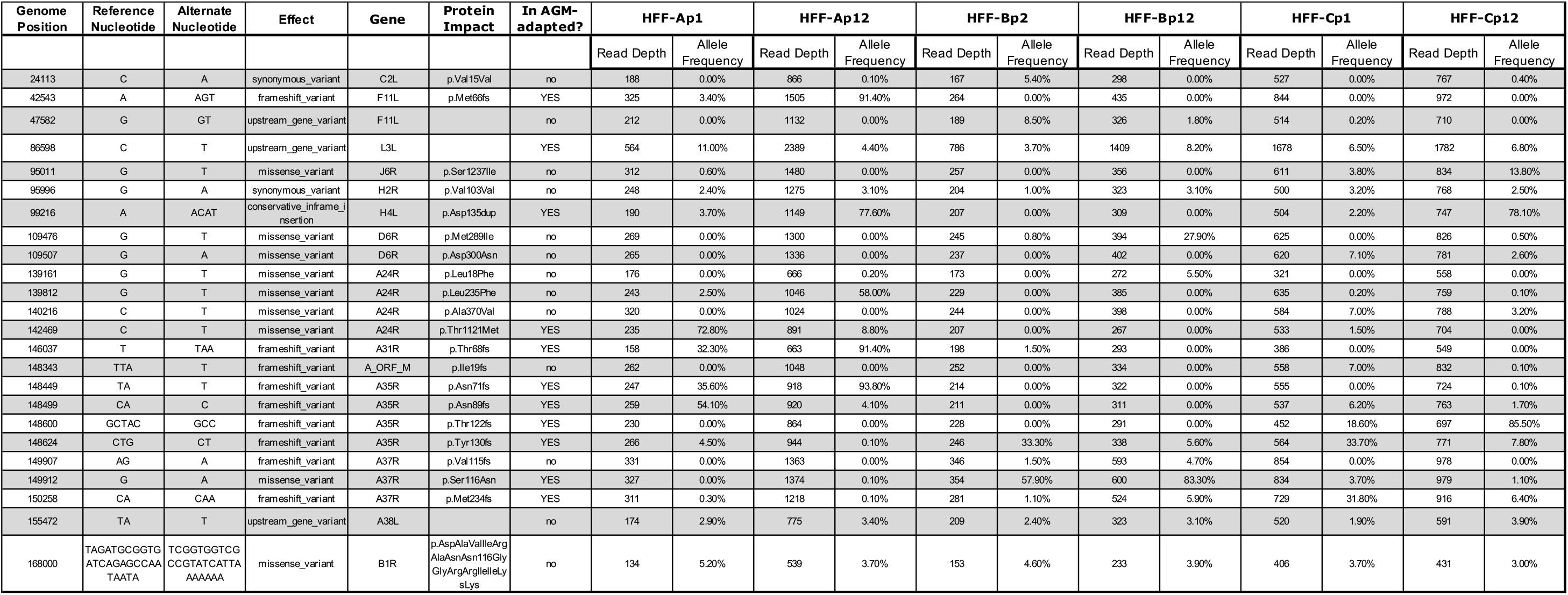
Nucleotide polymorphisms identified during experimental evolution of AGM-adapted VACVΔEΔK+RhTRS1 in human fibroblasts.

We were particularly interested in the fate of two AGM-adaptive SNPs (A24R* and A35R*) that were pre-existing in these populations and were individually sufficient to fully rescue VACVΔEΔK+RhTRS1 replication in AGM fibroblasts and partially rescue replication in HFF cells^16^. A24R* is a T1121M mutation in the catalytic subunit of the viral RNA polymerase (vRNAP) and A35R* is a TA211T frameshift in the immunomodulatory A35R gene predicted to truncate the 176 amino acid protein after 92 amino acids^30–32^. A24R* was only present in the founder population for HFF-A, initially at a 73% allelic frequency. This SNP rapidly declined in frequency, dropping to 19% frequency after only four passages, and to 9% allelic frequency at passage 12 (Fig. S5). All three founder populations had at least one of four different A35R frameshifts with an initial allelic frequency of at least 30% (Fig. S1). Populations HFF-A and HFF-C each initially contained two different frameshift variants, and during the course of the experiment the frequency of one allele increased in frequency while the other allele decreased in each population. HFF-B only had a single frameshift mutation which was lost during the course of the experiment.

Despite the loss of the T1121M mutation, we identified three SNPs in the A24R gene that were unique to HFF-adapted viruses: L18F, L235F, and A370V. Two of these, A370V and L18F, were only present in population HFF-B at an allelic frequency of approximately 5%, and thus are unlikely to represent adaptive mutations. However, L235F increased to an allelic frequency of >58% in HFF-A by passage 12. In addition to these A24R mutations, we also identified SNPs in several other transcription related genes. Populations HFF-A and HFF-C both contained a single amino acid insertion in H4L, which encodes the RNA polymerase-associated transcription-specificity factor RAP94. HFF-B contained a missense mutation in D6R, the helicase component of the vaccinia early transcription factor^33^ at 25% allelic frequency by passage 12. J6R, which encodes the large subunit of the vRNAP, had a S1237I SNP exclusive to HFF-C that increased to 14% allelic frequency by passage 12.

There were two additional pre-existing high frequency SNPs in the founder populations. An indel in A31R increased in frequency from 33% to 91% in HFF-A, and an S116N missense mutation in A37R increased from 51% to 83% in HFF-B. We also identified a frameshift in F11L, which alters cytoskeletal motility during infection^34^. This frameshift, present only in HFF-A, increased from 3% to 91% allelic frequency over the course of the experiment. Overall, many of these SNPs occur in essential genes that either are components of or are closely associated with the vRNAP. However, although multiple SNPs accumulated to high frequency during experimental evolution, no SNP was shared between all three populations, suggesting that there may be multiple pathways to HFF adaptation.

### HFF-adapted virus populations completely inhibit both PKR and RNase L pathways

To determine whether the process of adaptation to human cells enabled these viruses to inhibit both PKR and RNase L, we performed immunoblot analysis and RNA degradation assays on A549 cells infected with VACV-βg, VACVΔEΔK+RhTRS1, and passage 12 of each of the HFF-adapted populations. By immunoblot analysis each HFF-adapted population had low eIF2α phosphorylation levels, similar to VACV-βg levels, indicating that these viruses also inhibit the PKR pathway (Fig. 7A). In our previous studies, AGM-adapted viruses did not reduce PKR phosphorylation, only eIF2a phosphorylation was reduced^15,16^. However, in this experiment, cells infected with all three passaged populations had less PKR phosphorylation relative to VACVΔEΔK+RhTRS1 infected cells. This phenotype is different than the inhibition of the PKR pathway after PKR phosphorylation that we previously observed during adaptation in AGM cells^15,16^. However, it is in agreement with a recent report showing that duplication of RhTRS1 also reduced PKR phosphorylation in RhCMV infected human fibroblasts^21^. We observed similar trends in the degradation patterns of total RNA isolated from infected cells. While VACVΔEΔK+RhTRS1 again showed extensive RNA degradation, we did not detect RNA degradation products in cells infected with VACV-βg or any of the three HFF-adapted populations (Fig. 7B). Taken together, these data demonstrate that all three virus populations acquired the ability to fully inhibit human PKR and RNase L during experimental evolution.

**Figure 7.**
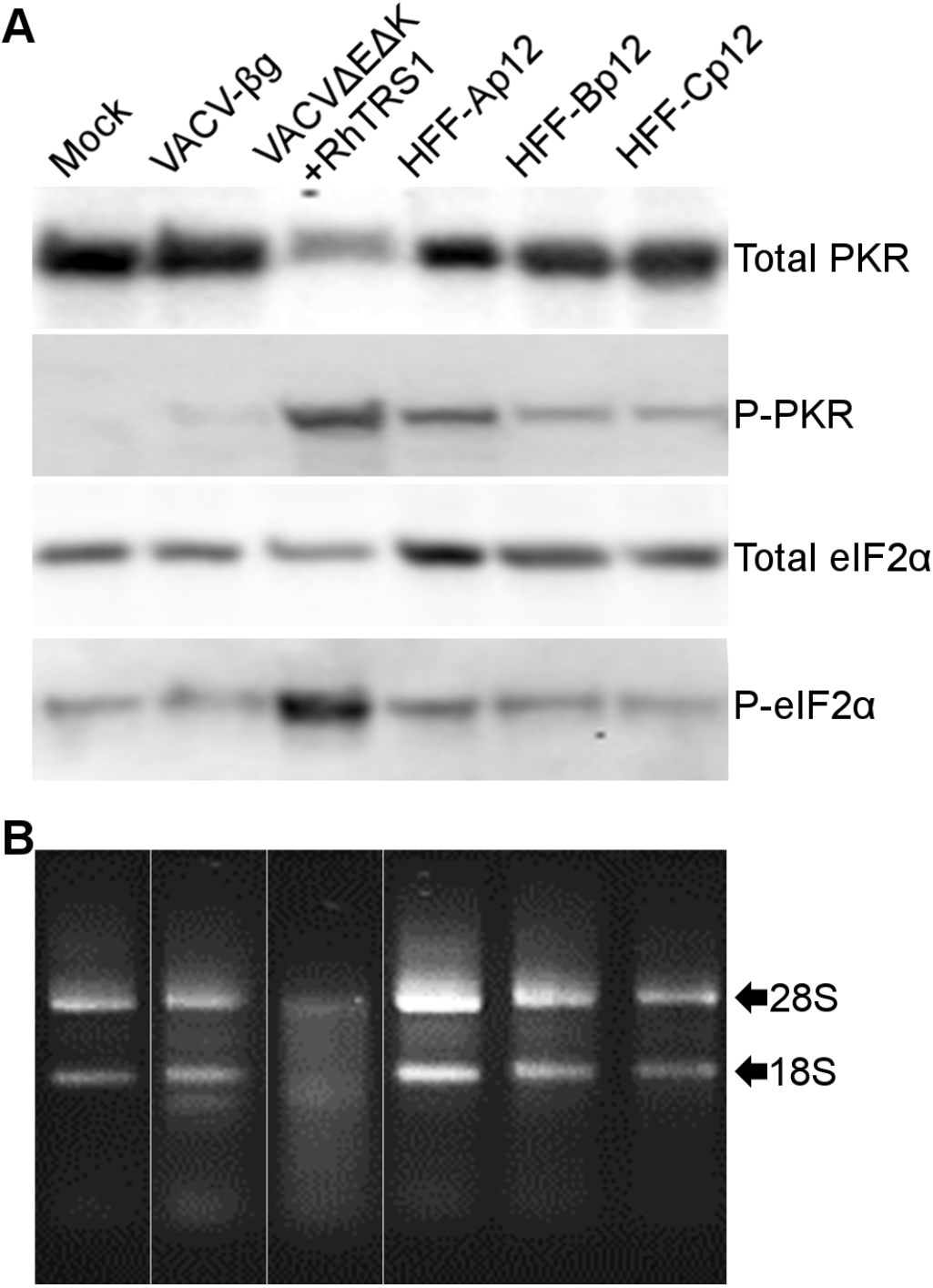
Passage 12 viruses inhibit both PKR and RNase L pathways in human cells. (A) A549 cells were infected with the indicated viruses (MOI = 3.0). One day post-infection, we collected either cell lysates (A) or total RNA (B). Cell lysates were analyzed by immunoblotting with the indicated antibodies (A). Total RNA was visualized on an agarose gel + 1% bleach (B). White bars separate lanes that were moved to align with panel A; however, all lanes shown were run on the same gel.

One possible explanation for the bimodal increase in replication fitness we observed during experimental evolution is that adaptation to PKR and RNase L may have occurred sequentially rather than simultaneously. To test this hypothesis and determine how PKR and RNase L susceptibility changed over the course of the experimental evolution, we analyzed HFF-Ap5 (immediately before the second fitness increase), HFF-Ap9 (immediately after the second fitness increase) and HFF-Ap12. Across these viruses, we identified a gradual decrease in phosphorylated PKR and eIF2α levels (Fig. 8A, upper panel). HFF-Ap12 infection showed the lowest amount of PKR phosphorylation, only slightly higher than VACV-βg infected cells. HFF-Ap5 infected cells had a decrease in eIF2α phosphorylation consistent with at least partial inhibition of the PKR pathway, while HFF-Ap9 and HFF-Ap12 had very little eIF2α phosphorylation, consistent with both populations replicating as well as VACV-βg.

**Figure 8.**
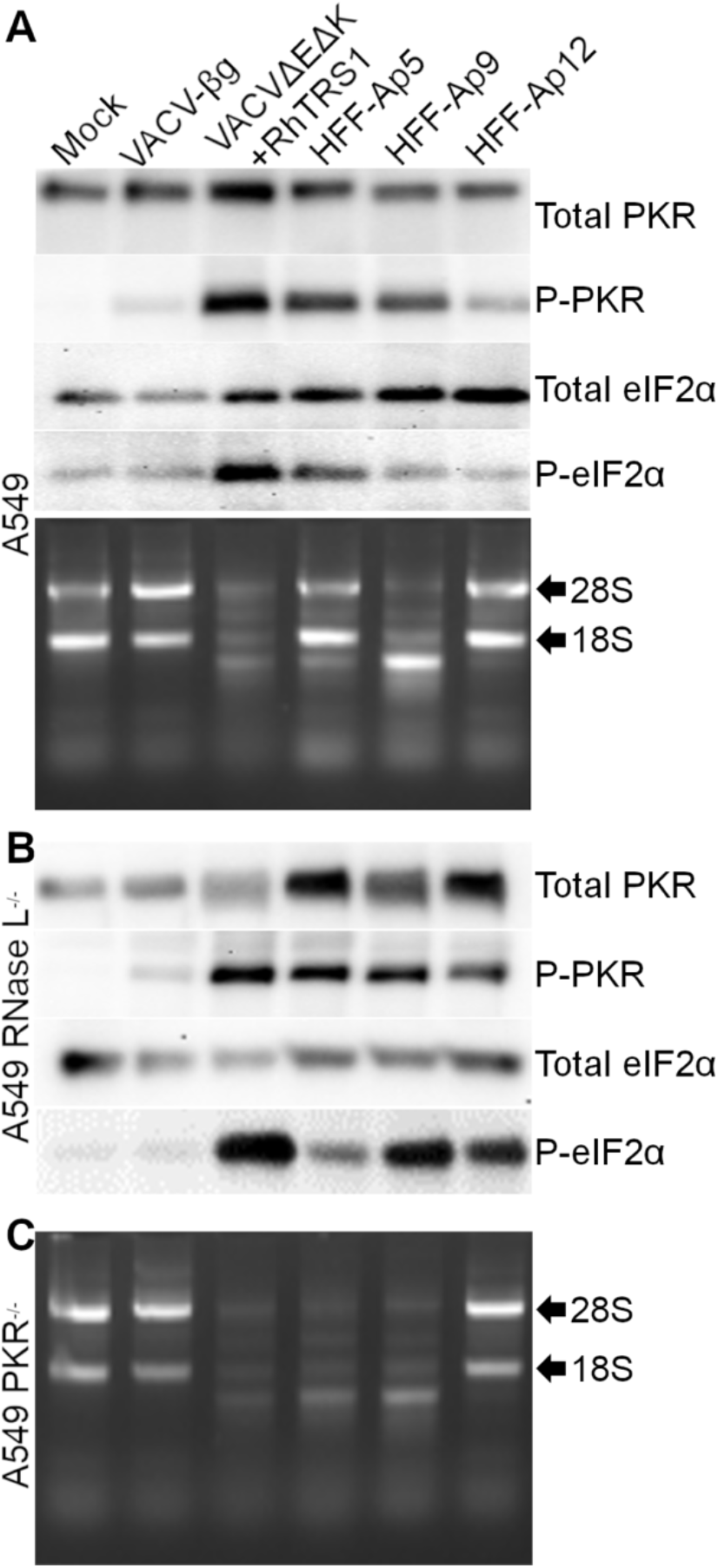
Phenotypic changes to PKR and RNase L inhibition occur even after replication is fully rescued in human cells. (A) A549 cells were infected with the indicated viruses (MOI = 3.0). One day post-infection, we collected either cell lysates (top panel) or total RNA (bottom panel). Cell lysates were analyzed by immunoblotting with the indicated antibodies. Total RNA was visualized on an agarose gel + 1% bleach. (B) A549 RNase L^-/-^ cells were infected with the indicated viruses (MOI = 3.0). One day post infection, cell lysates were analyzed by immunoblotting with the indicated antibodies. (C) A549 PKR^-/-^ cells were infected with the indicated viruses (MOI = 3.0). One day post infection, total RNA was harvested and visualized on an agarose gel + 1% bleach. <u>Data are representative of three independent biological replicates</u>.

**Figure 9.**
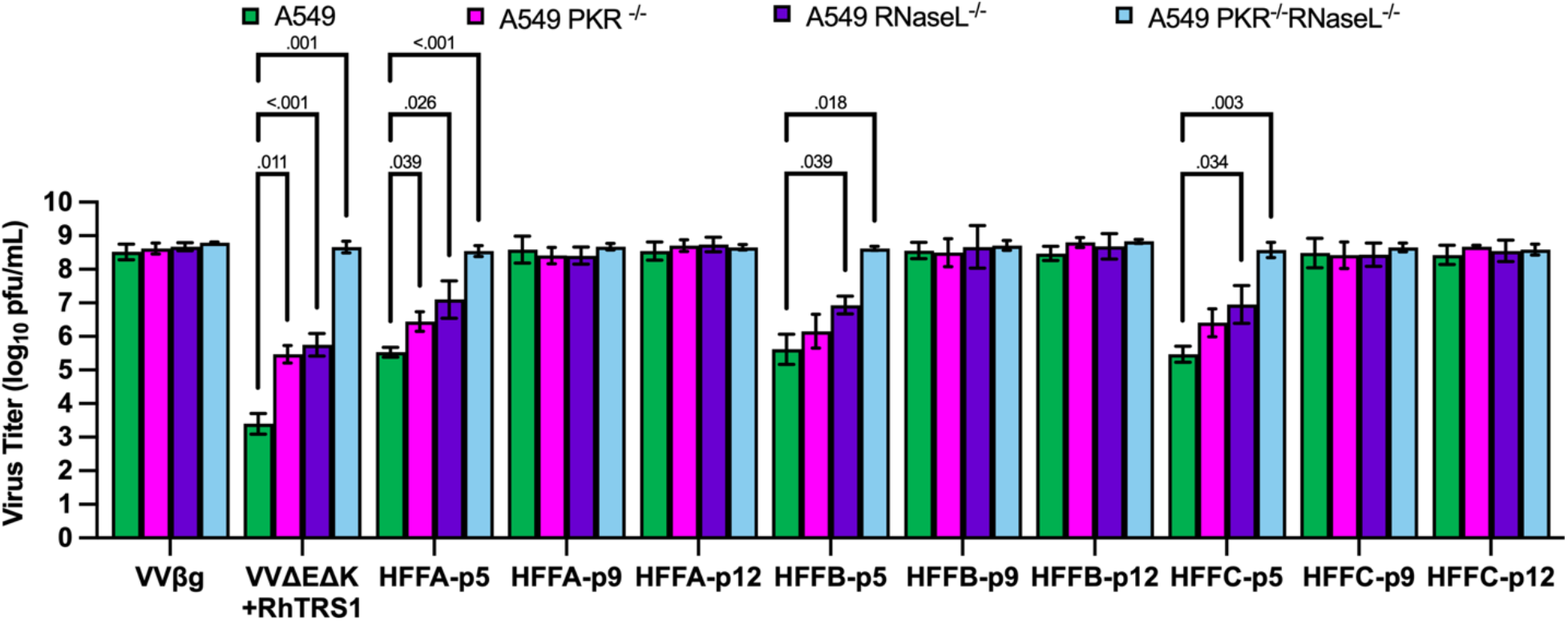
Population HFF-A adapted to PKR and RNase L concurrently. A549 cells (green bars), A549 PKR^-/-^ cells (magenta bars), A549 RNase L^-/-^ cells (purple bars), or A549 PKR^-/-^ RNase L^-/-^ cells (blue bars) were infected with the indicated viruses (MOI = 0.1). Two days post-infection, titers were determined by serial dilution on permissive BSC40 cells. <u>Columns represent the mean of three independent biological replicates</u>. Error bars indicate +/- one standard deviation. <u>Differences between samples were determined by two-way ANOVA. Samples with multiplicity adjusted p-values < 0.05 (Dunnett’s multiple comparison’s) are indicated by brackets with p-values indicated above</u>.

To determine if there were any differences in RNase L inhibition for these same virus populations, we infected A549 cells and performed an RNase L degradation assay (Fig. 8A, lower panel). HFF-Ap5 infected cells had a reduced level of RNA degradation products relative to VACVΔEΔK+RhTRS1 infected cells, consistent with partial but incomplete rescue of virus replication. Surprisingly, even though HFF-Ap9 replicated as well as VACV-βg in A549 cells, there was still substantial RNase L activity in these cells. However, HFF-Ap12 infected cells showed little to no evidence of RNA degradation, suggesting that there is a difference in the ability of HFF-Ap9 and HFF-Ap12 to fully inhibit RNase L activation even though both populations replicate to equivalent titers.

To rule out the possibility of a difference in replication efficiency between HFF-Ap9 and HFF-Ap12 in A549 cells that was not apparent in primary human fibroblasts, we measured viral titers in wildtype, PKR or RNase L single knockout, and double knockout A549 cells (Fig. 9). As in infected HFF cells, HFF-Ap5 replicated approximately 100-fold better than VACVΔEΔK+RhTRS1, and 100-fold less well than VACV-βg. This replication defect was partially improved by knocking out either PKR or RNase L, suggesting that both host restriction factors still exert at least some antiviral activity on HFF-Ap5. However, when both PKR and RNase L were knocked out, HFF-Ap5 replicated as well as VACV-βg. Both HFF-Ap9 and HFF-Ap12 replicated to similar titers, essentially as well as VACV-βg in all cell types tested consistent with their replication phenotype in HFF cells.

These data suggested that both restriction factors were gradually inhibited during adaptation, rather than sequentially inhibited. To test this hypothesis, we performed immunoblot analysis or RNA degradation assays on infected A549 RNase L^-/-^ cells or A549 PKR^-/-^ cells, respectively. In the absence of RNase L, all three HFF-A isolates resulted in PKR phosphorylation (Fig. 8B). There may be some reduction in PKR phosphorylation in the HFF-Ap12 infected cells; however, the reduction was consistently less than in infected wildtype A549 cells. As with wildtype A549 cell infection, we observed a partial reduction of eIF2α phosphorylation in HFF-Ap5 infected A549 RNase L^-/-^ cells. Although still lower than VACVΔEΔK+RhTRS1-infected cells, there was a relative increase in eIF2α phosphorylation in HFF-Ap9 or HFF-Ap12 infected cells, with an apparent peak at passage 9. Similarly, in the absence of PKR, we still observed substantial RNA degradation in HFF-p5 and HFF-p9 infected cells, but we did not observe RNA degradation in HFF-Ap12 infected cells (Fig. 8C). These data suggest that selective pressures were still driving phenotypic changes in both the PKR and RNase L response between passages 9 and 12 in all three populations even though the viruses at both timepoints replicated as well as VACV-βg. Taken together, this study suggests that initial adaptation by gene amplification acted as a “molecular foothold” to improve viral replication in otherwise resistant human cells, and thereby facilitate the emergence of novel, potentially species-specific adaptations that allow improved replication in the new host.

## Discussion

Intermediate hosts can increase contact between a virus and a new host species, and also drive adaptive changes, like gene amplification, that may improve the ability of a virus to replicate in these new species. Thus, the process of adaptation in one host may alter the likelihood of virus transmission to a variety of different species. We previously demonstrated that VACVΔEΔK+RhTRS1 failed to replicate in human fibroblasts to a level sufficient to sustain transmission upon serial passage. In this study we show that this increased resistance is due to human-derived cells having multiple blocks to VACVΔEΔK+RhTRS1 replication relative to AGM-derived cells. While adaptation in AGM fibroblasts was a critical intermediate step to expand the viral tropism, this adaptation did not fully inhibit either PKR or RNase L in human cells. however, during experimental evolution these viruses were able to adapt to human cells in a bimodal pattern. During this adaptation, some of the mutations in the founder populations that arose during AGM-adaptation were rapidly lost in human cells. It’s unclear if this rapid loss of some SNPs was due to bottlenecking effects of our experimental evolution strategy, or if these SNPs were in some way maladaptive in human cells. However, several mutations in the human adapted viruses dramatically increased in frequency, including some that were not present in the AGM-adapted viruses. Thus, after collapse of the *rhtrs1* locus, identifying the shared origin of the AGM- and human-adapted viruses may be obscured, emphasizing the need to identify early, transient biomarkers of rapid adaptation, such as gene duplication, that may indicate viruses poised to more easily cross species barriers.

Gene amplification is a well-established evolutionary mechanism, enabling organisms to rapidly respond and adapt to a given selective pressure. Examples of adaptation through gene amplification has been well documented in archaea, bacteria, and eukaryotes^35,36^. This phenomenon has more recently been recognized as a potent adaptive response in both poxviruses and herpesviruses^15,18–21^. In AGM cells, *rhtrs1* peaked at two copies per genome on average, and RhTRS1 overexpression alone was sufficient to fully rescue replication in AGM cells. We have previously shown that RhTRS1 overexpression alone is not sufficient to fully rescue VACVΔEΔK+RhTRS1 replication in human cells^15^. This current study confirms this observation that AGM-A expresses RhTRS1 higher than VACVΔEΔK+RhTRS1, yet is not fully rescued (Fig. S5). HFF-B and HFF-C each peaked at nearly double the average *rhtrs1* copy number reached in AGM cells, with 3.5 copies per genome (compare passage 6 in Fig. 4 and Fig. 5b), and substantially higher RhTRS1 protein expression than the AGM-A founder virus (Fig. S4), yet they still did not replicate as well as VACV-βg. In contrast, HFF-Ap5 expressed very little RhTRS1 despite further amplification of the locus relative to the founder AGM-A population, while HFF-Ap9 and HFF-Ap12 both express substantially more RhTRS1 than HFF-Ap5, and as much or more than any timepoint in HFF-B or HFF-C, despite having lower average *rhtrs1* copy numbers (Figs. 6 and S4). We do not have a compelling explanation for this lack of expression at an early timepoint. Notably, passage 5 is the timepoint that shows a stereotypical decrease in virus titer which has been observed in multiple poxvirus experimental evolution studies^15,19^. In this instance the dip in virus replication may somehow be associated with the low RhTRS1 expression. Taken together, these observations may suggest that the dual inhibition by both PKR and RNase L in human cells is not as susceptible to evasion by *rhtrs1*. This hypothesis is supported by our observation that PKR phosphorylation and RNA degradation were both decreased but not eliminated at passage 5 (Fig. 6 a and b). Alternatively, it’s possible that this amplification marks the limits of the VACV genome’s capability to accommodate additional genetic material, rather than an inherent inability to inhibit both restriction factors given sufficient overexpression. Nevertheless, although *rhtrs1* amplification alone was insufficient to fully rescue virus replication, our study demonstrates that pre-existing copy number variation can both expand viral host tropism and facilitate very rapid responses to new selective pressures.

Consistent with our previous results, *rhtrs1* amplification and the corresponding overexpression of RhTRS1 only partially inhibited PKR activation in human cells. This observation is surprising in light of a recent study demonstrating that rhesus cytomegalovirus was able to productively infect human cells, at least in part due to an overexpression of RhTRS1 relative to human cytomegalovirus encoded orthologs^37^. Although this relative overexpression of RhTRS1 permitted replication in human fibroblasts, PKR knockout still improved virus replication. Furthermore, in this same study RhCMV serial passage in human cells resulted in a duplication of *rhtrs1*^21^. Unlike our study, in the context of RhCMV this duplication fully rescued replication in human cells. Taken together, these differences in PKR antagonism between RhTRS1 expressed by RhCMV and RhTRS1 expressed by the chimera VACVΔEΔK+RhTRS1, or other VACV-based assays^21,23,37^ suggest that differences in the intracellular environment induced by infection with these two viruses may influence activation of different host restriction factors.

Knocking out either PKR or RNase L individually provided comparable improvement in VACVΔEΔK+RhTRS1 replication fitness (Figs. 2, 3, and 8). Complete rescue of viral replication only occurred in the absence of both PKR and RNase L (Figs. 4 and 8). Moreover, knocking out either restriction factor individually resulted in a very similar partial increase to VACVΔEΔK+RhTRS1 replication, similar to AGM-adapted viruses. These observations suggest that both antiviral responses act as independent barriers to virus replication.

Although infection with the AGM-adapted viruses all resulted in PKR activation, PKR knockout did not result in any increase in virus titer. One possibility is that these viruses dampen but do not eliminate PKR activation. Therefore, the viruses might be able to replicate well early, and it may be that sufficient dsRNA to activate PKR is only present later in the infection when it is too late to have a detectable impact on viral replication. This hypothesis is supported by a previous studied showed VACV lacking the amino-terminus of the PKR antagonist E3L (VACVΔ83N) replicates as well as wildtype VACV in HeLa cells with or without PKR. However, at late timepoints this virus also activates PKR without significantly impacting the titer^38,39^.

We did identify a small but statistically significant increase in one of the AGM-adapted populations (AGM-A), in RNase L^-/-^ cells (Fig 3). Small cleavage products produced by active RNase L have been shown to dimerize and act as additional PKR substrates^40,41^. Therefore, the modest increase in AGM-A replication in the absence of RNase L might indicate that crosstalk between the PKR and RNase L pathways also plays a minor role in restricting AGM-adapted virus replication in human cells. We have previously demonstrated that RNase L is functional in AGM fibroblasts, and thus inactivation cannot explain the difference in RNase L activation between cells from the two species. Therefore, the most likely explanation is some difference in the course of the VACVΔEΔK+RhTRS1 infection that favors RNase L activation only in the human cells. One possibility may be differences in dsRNA production between the two cell types, as has been shown in in other poxvirus systems^42,43^. Alternatively, at the host level, AGM OAS3 is predicted to have one less double-stranded RNA binding domain than human OAS3, which may alter the activation threshold for 2-5A production. Furthermore, there are 54 aa differences between human and AGM RNase L, including regions implicated in determining the rate of RNA cleavage^44^ and in 2-5A binding^45^ (Fig. S6). These host genetic differences in RNase L may contribute to the phenotypic differences we observed in response to VACVΔEΔK+RhTRS1 infection.

Apart from the species-specific differences in host restriction factors, we also observed species-specific differences in the kinetics of virus adaptation to human cells as compared to AGM cells. The virus evolved bimodally in HFFs, with an initial rapid 10-fold increase in replication, followed by a somewhat slower second increase in viral replication. Despite the bimodal curve of this adaptive profile the restriction factors were not inhibited sequentially. Instead, there was a gradual decrease in both PKR activation and RNase L activation throughout the entire adaptive process (Fig. S5). Furthermore, even after the three different populations were able to replicate as efficiently as VACV-βg by passage 9, we observed continued phenotypic changes in both PKR and RNase L inhibition at passage 12 (Fig. 8). In A549 cells, RNase L activity is more pronounced at passage 9 than at either passage 5 or 12 (Fig. 8A, bottom panel). One possible explanation for this phenotype is that because PKR is better inhibited at passage 9 than at passage 5 (Fig. 8A, top panel) virus replication is improved. As a consequence, dsRNA accumulation may increase because more replicating virus is present, ultimately stimulating more RNase L activity as the virus populations adapted to human cells. Previous reports in both CMV and VACV-based systems have shown RhTRS1 inhibiting the PKR pathway at a stage after PKR phosphorylation but before eIF2α phosphorylation. However, in this study, all human cell adapted viruses inhibited PKR phosphorylation substantially better at passage 12 than either the founder AGM-adapted viruses, or earlier human passages of this virus (Fig. 7 and Fig. 8A). This reduction in PKR phosphorylation is consistent with a recent report demonstrating that RhTRS1 duplication blocked PKR phosphorylation in RhCMV infected HFFs. These differences in PKR phosphorylation may be a result of overexpression resulting from the increased copy number relative to AGM-adapted viruses in this study, and duplication of *rhtrs1* in the RhCMV study. It may also be that the evolved SNPs outside the *rhtrs1* locus evade PKR activation independent of RhTRS1. For example, the mutations we identified in various components of the viral transcriptional machinery may alter transcriptional kinetics, possibly reducing dsRNA production below the activation threshold of these host restriction factors. These experiments are currently ongoing.

As with our previous experiments adapting VACVΔEΔK+RhTRS1 in AGM cells, we did not identify a single SNP in *rhtrs1*. Since that initial study, the VACV decapping enzymes D9 and D10 have each been shown to inhibit both PKR and RNase L activity. These enzymes act in concert with the host exoribonuclease Xrn1 to deplete both host and viral RNA, thereby reducing total intracellular dsRNA^22,46–48^. However, as with *rhtrs1*, we did not detect any mutations in either of these VACV genes. It’s therefore unclear whether these genes play any role in adaptation of this chimeric virus. Because we passaged viruses that had already been adapted to AGM fibroblasts, some virus populations contained the A24R* (T1121M) and A35R* (TA211T indel) mutations that were individually sufficient to fully rescue VACVΔEΔK+RhTRS1 in AGM cells and partially rescue replication in human fibroblasts^16^. It should be noted that only the AGM-A founder population included these two SNPs, yet all 3 populations underwent the same bimodal adaptation, suggesting that *rhtrs1* copy number variation alone was sufficient for the initial adaptation in human cells. Furthermore, despite the replication benefit conferred by A24R* in isolation, the frequency of this variant in the population plummeted from approximately 78% to less than 20% after only four rounds of serial passage, suggesting that there may not be a straight-forward evolutionary path for the T1121M mutation to further adapt to inhibit human PKR, or that it may in some way be maladaptive in human cells.

Although the T1121M variant was lost, two new variants emerged in A24R during serial passage in human cells. L18F emerged in population HFF-B, and has been reported previously during serial passage of VACVΔE3L in HeLa cells^49,50^. However, in that study it only provided a 3-fold increase to virus replication, and paradoxically increased PKR activation in these cells. Thus, L18F is unlikely to explain the increase in replication fitness in these primary human fibroblasts. The third A24R mutation we identified, L235F, emerged in population HFF-A and maps to a conserved residue in the alpha-5 helix of the “lobe” domain of A24R (rpo132). This domain, together with the “clamp head” domain of J6R (rpo147) guides the DNA deeper into the holoenzyme to the site of transcription bubble formation^51,52^. We also identified mutations in other components of the vRNAP. A D135DD insertion in the H4L gene was present in both populations HFF-A and HFF-C. H4L encodes the RAP94 protein, a poxvirus-specific transcription factor that has no known homolog in other species^51^. H4L is expressed late during virus replication, but only associates with the vRNAP during early transcription. Previous studies have implicated H4L in both the recognition of early replication stage viral promoters^53^ as well as having a role in the elongation and precise termination of these transcripts^54^ through its direct interactions with NPH I, VETF, and other components of the holoenzyme^54,55^. H4L is also necessary to efficiently package the RNA polymerase and other components of the VACV transcriptional apparatus during virion assembly^56^. Another SNP unique to these human passaged viruses occurred in D6R, which encodes the viral early transcription factor (vETFs). This variant was only present in population HFF-B which lacked the H4L SNP. Taken together, although none of these SNPs were shared in all three human passaged virus populations, each population had a SNP in a gene involved in early transcription initiation and termination. The accumulation of multiple mutations in the vRNAP suggests the potential for a common adaptive pathway, and the different array of SNPs that accumulated between the three populations may represent a balance between evasion of PKR/RNase L activity, and maintaining the critical interactions between the various subunits of the vRNAP. Taken together, these results suggest that altering vRNAP function and transcription might play an important role in poxvirus adaptation.

Combined with our previous work adapting this virus to African green monkey cells, these experiments suggest one possible model for initial spillover events. Individuals within a population may have a spectrum of susceptibilities, similar to the differences in VACVΔEΔK+RhTRS1 susceptibility we reported for different AGM-derived cell lines^15,16^.

These differences are likely driven in part by host restriction factor variation influencing the threshold necessary for a virus to overcome restriction and productively infect a new host^6^. In a population with more variability in restriction factor antiviral activity, susceptible individuals provide opportunities for the virus to continue to circulate, and more resistant individuals provide selective pressure, facilitating emergence of variants including gene duplication. Thus, these populations may be more prone to drive cross-species transmission, in a manner similar to our data demonstrating that adaptation in AGM fibroblasts was necessary to provide a “molecular foothold” for the virus to subsequently adapt to human cells. Currently, however, little is known about intraspecies variation in host immune responses, beyond some compelling examples such as the Mamu-A*01 MHC allele in rhesus macaques that attenuates disease progression in SIV-infected rhesus macaques^57^. Identifying these populations with differential susceptibility may be critical to detecting emerging viruses early. Overall, our data supports our hypothesis that initial adaptation by gene amplification acts as a “molecular foothold” to broadly improve viral replication in resistant host species, and thereby facilitate the emergence of novel, potentially species-specific adaptations to maintain replication in the new host.

## Acknowledgements

We thank Bernard Moss (NIH), Denise Galloway (Fred Hutchinson Cancer Center), and Stan Riddell (Fred Hutchinson Cancer Center) for reagents. We thank Bala Burugula for assistance with sequencing library preparation. We thank members of the Rothenburg and Kitzman labs for helpful discussions and critical reading of the manuscript.

## Data availability

The Illumina sequencing dataset will be available on the Short Read Archive (SRA# PRJNA846067). Biological replicates not shown in this manuscript are available on the public repository Dryad. All other data is available in the manuscript.

## Author Contributions

Conceived and designed the experiments: GB, JOK, SB, CS, APG

Performed the experiments: SB, CS, GB

Analyzed the data: SB, CS, GB, JOK, SR

Wrote the paper: SB, CS, GB, JOK, SR, APG

---

Table S1. Structural variants detected in each population and associated metrics.

Table S2. Point and short insertion/deletion variants detected in each population and associated metrics.

