## Supplementary material for "Gene amplification acts as a molecular foothold to facilitate cross-species adaptation and evasion of multiple antiviral pathways": Figure S1

| | 1 | | | | 50 |
|---|---|---|---|---|---|
| A35R_WT | MDAAFVITPM | GVLTITDTLY | DDLDISIMDF | IGPYIIGNIK | TVQIDVRDIK |
| **A35R_Asn71fs** | MDAAFVITPM | GVLTITDTLY | DDLDISIMDF | IGPYIIGNIK | TVQIDVRDIK |
| A35R_Asn89fs | MDAAFVITPM | GVLTITDTLY | DDLDISIMDF | IGPYIIGNIK | TVQIDVRDIK |
| **A35R_Thr122fs** | MDAAFVITPM | GVLTITDTLY | DDLDISIMDF | IGPYIIGNIK | TVQIDVRDIK |
| A35R_Tyr130fs | MDAAFVITPM | GVLTITDTLY | DDLDISIMDF | IGPYIIGNIK | TVQIDVRDIK |
| A35R_Asn159fs | MDAAFVITPM | GVLTITDTLY | DDLDISIMDF | IGPYIIGNIK | TVQIDVRDIK |

| | 51 | | | | 100 |
|---|---|---|---|---|---|
| A35R_WT | YSDMQKCYFS | YKGKIVPQDS | NDLARFNIYS | ICAAYRSKNT | IIIACDYDIM |
| **A35R_Asn71fs** | YSDMQKCYFS | YKGKIVPQDS | *MIWLDSTFIA* | *FVPHTDQKIP* | *SS*-------- |
| A35R_Asn89fs | YSDMQKCYFS | YKGKIVPQDS | NDLARFNIYS | ICAAYRS*KIP* | *SS*-------- |
| **A35R_Thr122fs** | YSDMQKCYFS | YKGKIVPQDS | NDLARFNIYS | ICAAYRSKNT | IIIACDYDIM |
| A35R_Tyr130fs | YSDMQKCYFS | YKGKIVPQDS | NDLARFNIYS | ICAAYRSKNT | IIIACDYDIM |
| A35R_Asn159fs | YSDMQKCYFS | YKGKIVPQDS | NDLARFNIYS | ICAAYRSKNT | IIIACDYDIM |

| | 101 | | | | 150 |
|---|---|---|---|---|---|
| A35R_WT | LDIEDKHQPF | YLFPSIDVFN | ATIIEAYNLY | TAGDYHLIIN | PSDNLKMKLS |
| **A35R_Asn71fs** | ---------- | ---------- | ---------- | ---------- | ---------- |
| A35R_Asn89fs | ---------- | ---------- | ---------- | ---------- | ---------- |
| **A35R_Thr122fs** | LDIEDKHQPF | YLFPSIDVFN | A*NHRSV*---- | ---------- | ---------- |
| A35R_Tyr130fs | LDIEDKHQPF | YLFPSIDVFN | ATIIEAYNL*I* | *QLEIII*---- | ---------- |
| A35R_Asn159fs | LDIEDKHQPF | YLFPSIDVFN | ATIIEAYNLY | TAGDYHLIIN | PSDNLKMKLS |

| | 151 | | 176 |
|---|---|---|---|
| A35R_WT | FNSSFCISDG | NGWIIIDGKC | NSNFLS |
| **A35R_Asn71fs** | ---------- | ---------- | ------ |
| A35R_Asn89fs | ---------- | ---------- | ------ |
| **A35R_Thr122fs** | ---------- | ---------- | ------ |
| A35R_Tyr130fs | ---------- | ---------- | ------ |
| A35R_Asn159fs | FNSSFCIS*AA* | *MDGL*------ | ------ |

Figure S1. Amino acid alignment of different A35R predicted mutants. Amino acid alignment of the predicted VACV Copenhagen A35R gene product with the predicted A35R truncation mutants resulting from the frameshift mutations (Asn71fs, Asn89fs, Thr122fs, Tyr130fs, Asp159fs**)** identified in HFF-A, HFF-B and HFF-C virus populations. Amino acid differences introduced by the frame shift before the end of the truncated products are underlined and italicized. The Asn71fs (red) and Thr122fs (green) were present at allele frequencies >85% in virus populations HFF-A and HFF-C respectively.
