## Supplementary material for "Gene amplification acts as a molecular foothold to facilitate cross-species adaptation and evasion of multiple antiviral pathways": Figure S2

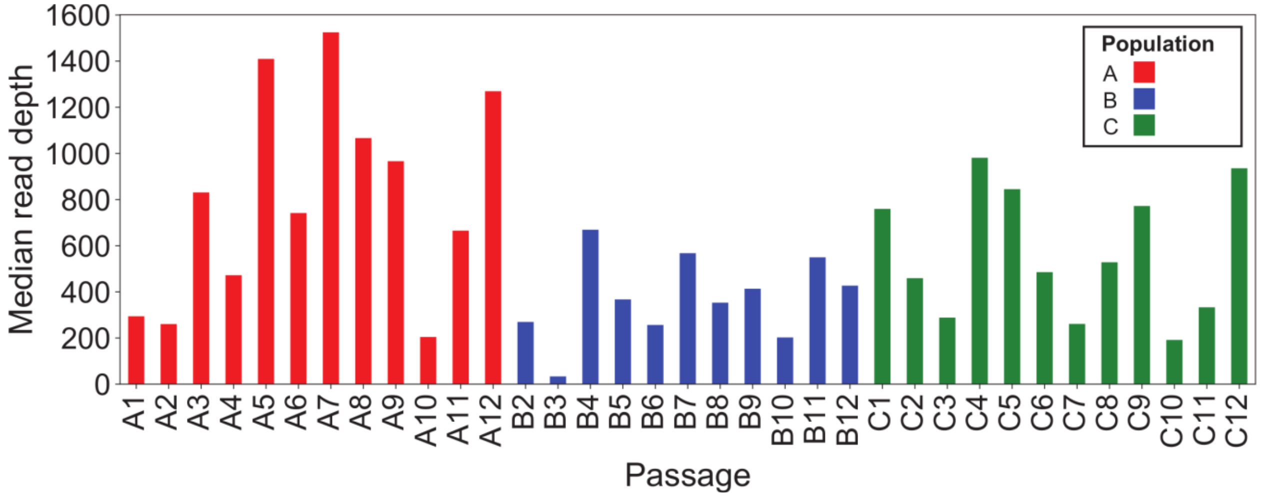


Figure S2. Median sequencing depth for each passage of populations A (red), B (blue), and C (green). The depth at each position was computed using bedtools, genomecov, and the median depth across all positions is shown on the y-axis.
