## Supplementary material for "Gene amplification acts as a molecular foothold to facilitate cross-species adaptation and evasion of multiple antiviral pathways": Figure S3

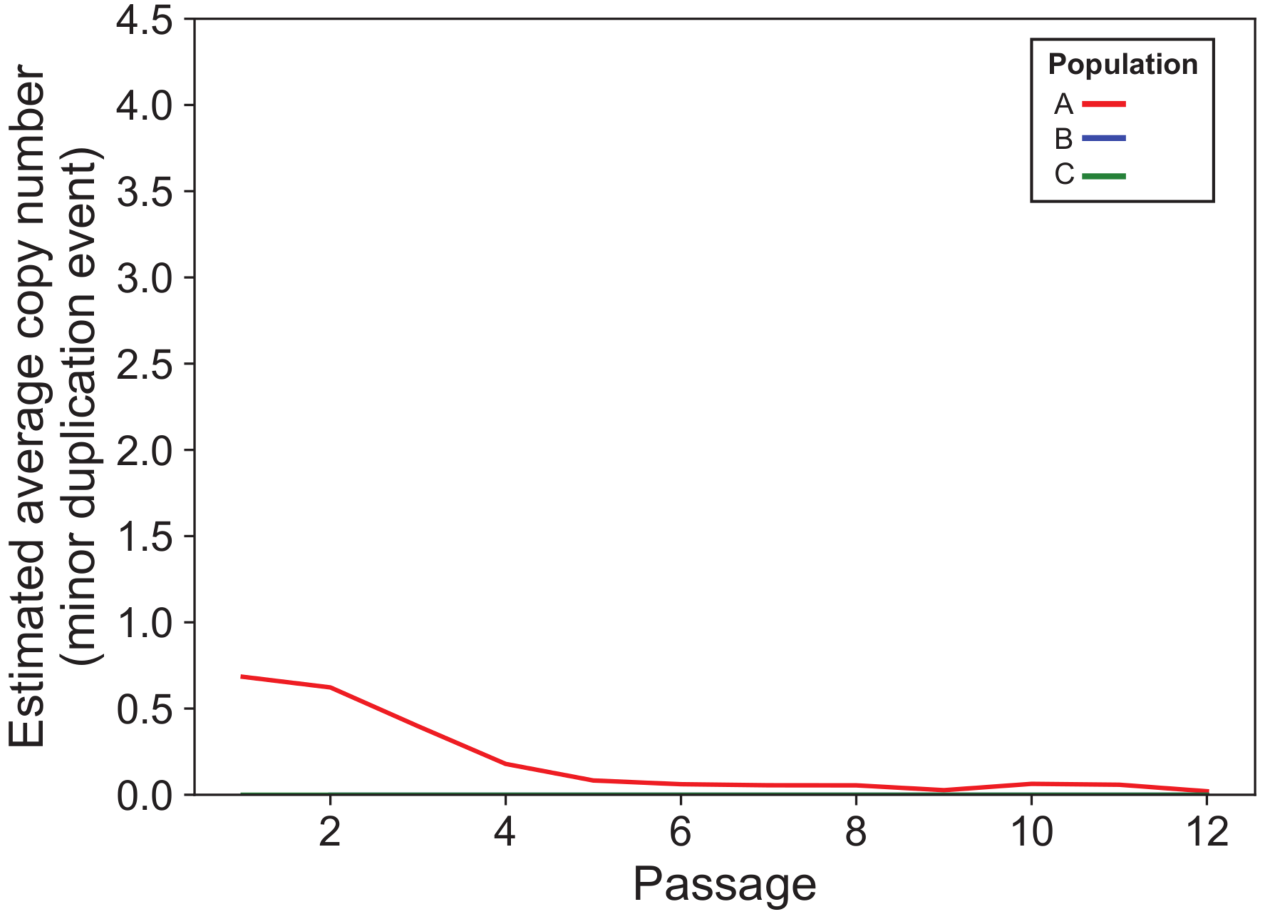


Figure S3. Estimated average copy number of the minor 3.4 kb duplication event spanning the NeoR gene to J2R by passage for the A (red), B (blue), and C (green) populations. Populations B and C are effectively coincident with the X-axis. Since the minor duplication event overlaps the major duplication event and both contribute to the read depth signal, we instead estimated the copy number by training a linear model to predict the EGFP adjusted read depth for the major duplication event from the frequency of breakpoint-containing split reads, and used it to predict the copy number contribution specific to the shorter duplication.
