## Supplementary material for "Gene amplification acts as a molecular foothold to facilitate cross-species adaptation and evasion of multiple antiviral pathways": Figure S4

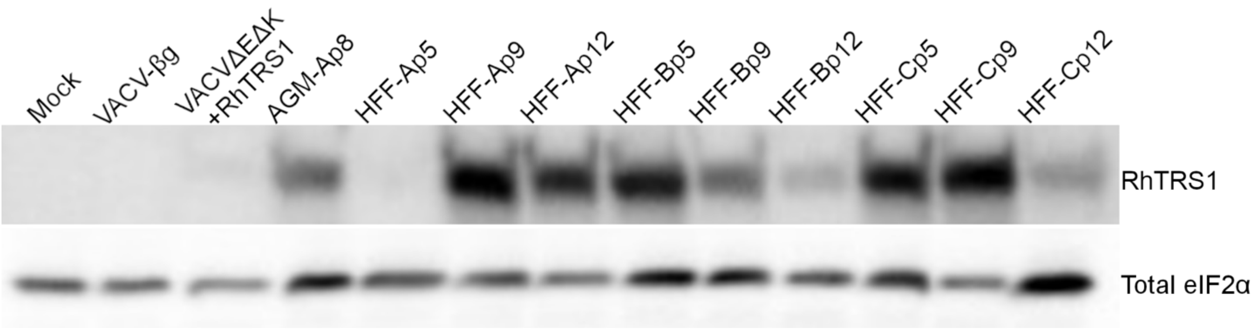


Figure S4. RhTRS1 expression in human cells. A549 cells were infected with the indicated viruses (MOI = 3.0). One day post infection, cell lysates were analyzed by immunoblotting with the indicated antibodies. Due to antibody availability, total eIF2α is used as a loading control in this experiment. Data are representative of two independent biological replicates.
