## Supplementary material for "Gene amplification acts as a molecular foothold to facilitate cross-species adaptation and evasion of multiple antiviral pathways": Figure S5

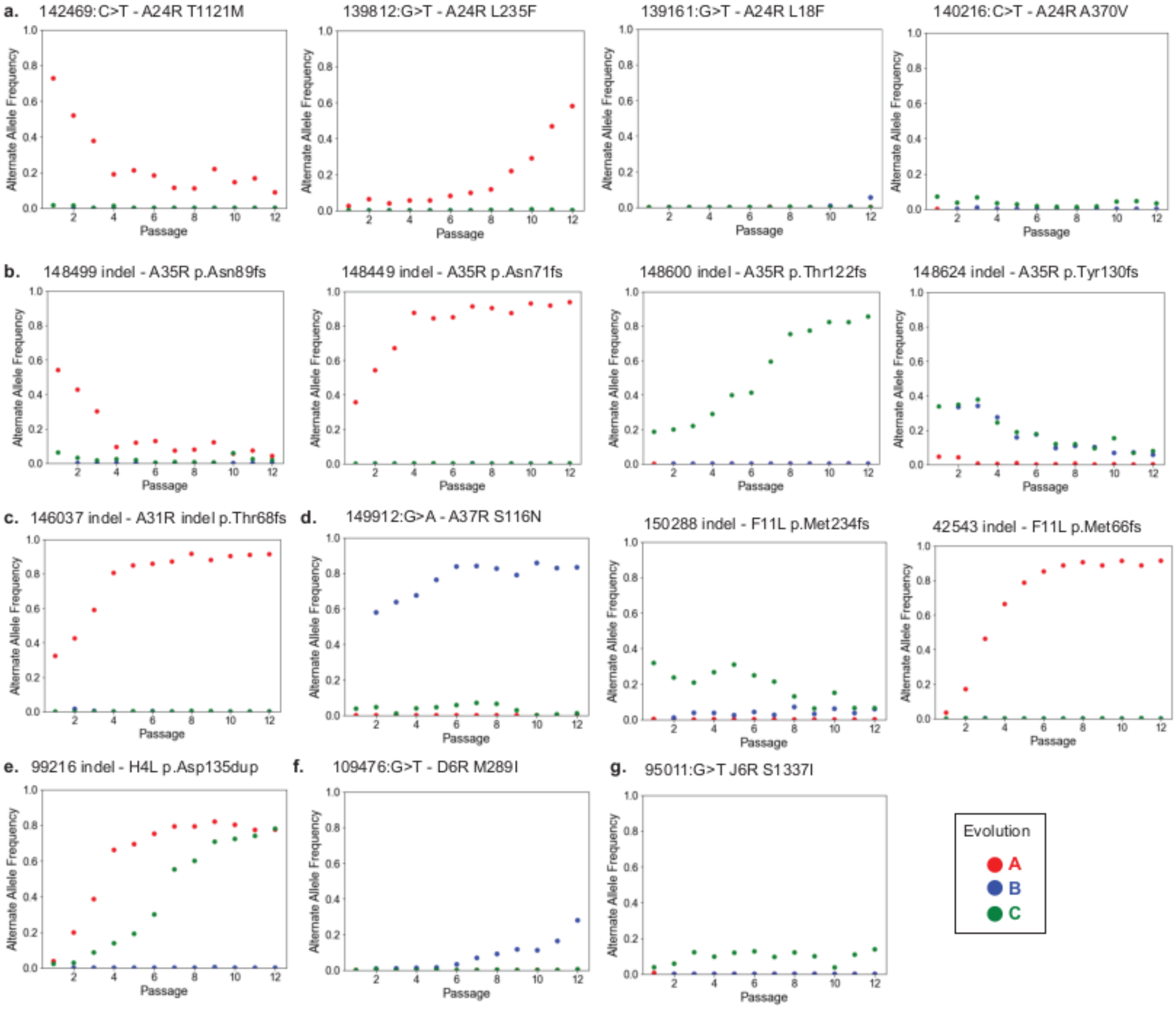


Figure S5. Dynamics of point and short indel mutations during HFF adaptation. Shown are variant allele frequencies (VAFs; y-axis), plotted against passage number, with each panel corresponding to a single mutation.
