## Supplementary material for "Gene amplification acts as a molecular foothold to facilitate cross-species adaptation and evasion of multiple antiviral pathways": Figure S6

```
                 1                                                     50
NP_066956_1      MESRDHNNPQ EGPTSSSGRR AAVEDNHLLI KAVQNEDVDL VQQLLEGGAN
XP_007987466_2   METRDHNNPQ EGPTSSSGRR ATVEDNHLLI KAVQTEDVDL VQQLLEGGAN
                 51                                                   100
NP_066956_1      VNFQEEEGGW TPLHNAVQMS REDIVELLLR HGADPVLRKK NGATPFILAA
XP_007987466_2   VNFQEKEGGW TPLHNAVQMD REDIVELLLR YDADPVLRKK NGATPFIVAA
                 101                                                  150
NP_066956_1      IAGSVKLLKL FLSKGADVNE CDFYGFTAFM EAAVYGKVKA LKFLYKRGAN
XP_007987466_2   IVGNVKLLEL FLSKGADVNE CDFYGFTAFM EAAVYGNVKA LKFLYERGAN
                 151                                                  200
NP_066956_1      VNLRRKTKED QERLRKGGAT ALMDAAEKGH VEVLKILLDE MGADVNACDN
XP_007987466_2   VNLRRKTKED QERLRKGGAT ALMDAAKEGH VEVLKILLDE MGADVNACDN
                 201                                                  250
NP_066956_1      MGRNALIHAL LSSDDSDVEA ITHLLLDHGA DVNVRGERGK TPLILAVEKK
XP_007987466_2   MGRNALIHAL LSSHNRDVEA ITHLLLDHGA DVNVRGERGK TPLILAVENK
                 251                                                  300
NP_066956_1      HLGLVQRLLE QEHIEINDTD SDGKTALLLA VELKLKKIAE LLCKRGASTD
XP_007987466_2   HLGLVKRLLK QEHIEINDTD SDGKTALLLA VELKLKDIAQ LLCDCGASTD
                 301                                                  350
NP_066956_1      CGDLVMTARR NYDHSLVKVL LSHGAKEDFH PPAEDWKPQS SHWGAALKDL
XP_007987466_2   CGDLVMIARR NYDHSLVKLL LSHGAEEHFH PPAEDWKPQS SHWGAALKDL
                 351                                                  400
NP_066956_1      HRIYRPMIGK LKFFIDEKYK IADTSEGGIY LGFYEKQEVA VKTFCEGSPR
XP_007987466_2   HRIYRPMIGK LKFFIDEKYK IADTSEGGIY LGFYEKQEVA VKTFCEGSPR
                 401                                                  450
NP_066956_1      AQREVSCLQS SRENSHLVTF YGSESHRGHL FVCVTLCEQT LEACLDVHRG
XP_007987466_2   ARQEVSCLQS SRENSHLVTF YGSESHRGHL FVCVTLCEQT LEACLEVHRG
                 451                                                  500
NP_066956_1      EDVENEEDEF ARNVLSSIFK AVQELHLSCG YTHQDLQPQN ILIDSKKAAH
XP_007987466_2   EDVENEEDEF ARNVLSSIFK AVQELHLSCA YTHQDLQPQN ILIDSKNAVH
                 501                                                  550
NP_066956_1      LADFDKSIKW AGDPQEVKRD LEDLGRLVLY VVKKGSISFE DLKAQSNEEV
XP_007987466_2   LADFDKSIKW TGDPQEVKRD LEDLGRLVLY VVKKGSISFE ELKAQSNEEV
                 551                                                  600
NP_066956_1      VQLSPDEETK DLIHRLFHPG EHVRDCLSDL LGHPFFWTWE SRYRTLRNVG
XP_007987466_2   VQLSPDEETK DLIHHLFHPG EHVRDCLGDL LGHPFFWTWE SRYRTLRNVG

                601                                                  650
NP_066956_1      NESDIKTRKS ESEILRLLQP GPSEHSKSFD KWTTKINECV MKKMNKFYEK
XP_007987466_2   NESDIKTRKH KSEILKLLQP GPSEHSVSFD KWTTKINADV MKKMNEFYKK

                 651                                                  700
NP_066956_1      RGNFYQNTVG DLLKFIRNLG EHIDEEKHKK MKLKIGDPSL YFQKTFPDLV
XP_007987466_2   SGNFYQNTVG DLLKFIRNLG EHIDEEKHKK MKLKIGDPSR YFQKTFPDLV
                 701                                         741
NP_066956_1      IYVYTKLQNT EYRKHFPQTH SPNKPQCDGA GGASGLASPG C
XP_007987466_2   IYVYTKLQNT EYRKHFPQTH SSNKPQCDGA GGTSRLASPG C
```

Figure S6. Amino acid alignment of African green monkey and human RNase L. Predicted RNase L amino acid sequences from AGM (XP_007987466.2) and human (NP_066956.1) RNase L were aligned using Clustal Omega. The two sequences share 93% identity. Amino acid differences in the ankyrin, protein kinase homology and ribonuclease domains are highlighted in red.
